# Combining RAS^G12C^(ON) inhibitor with SHP2 inhibition sensitises immune excluded lung tumours to immune checkpoint blockade: a strategy for turning cold tumours hot

**DOI:** 10.1101/2024.01.15.575765

**Authors:** Panayiotis Anastasiou, Christopher Moore, Sareena Rana, Andrea de Castro, Mona Tomaschko, Jesse Boumelha, Edurne Mugarza, Cristina Blaj, Sophie de Carné, Robert Goldstone, Jacqueline A.M. Smith, Elsa Quintana, Miriam Molina-Arcas, Julian Downward

## Abstract

Mutant selective drugs targeting the inactive, GDP-bound form of KRAS^G12C^ have been approved for use in lung cancer, but responses are short-lived due to rapid development of resistance. In this study we use a novel covalent tri-complex inhibitor, RMC-4998, that targets RAS^G12C^ in its active, GTP-bound form to investigate treatment of KRAS mutant lung cancer in various immune competent mouse models. While this RAS^G12C^(ON) inhibitor was more potent than the KRAS^G12C^(OFF) inhibitor adagrasib, rapid pathway reactivation was still observed. This could be delayed using combined treatment with a SHP2 inhibitor, RMC-4550, which not only impacted RAS pathway signalling within the tumour cells but also remodelled the tumour microenvironment (TME) to be less immunosuppressive and promoted interferon responses. In an inflamed, “hot”, mouse model of lung cancer, RAS^G12C^(ON) and SHP2 inhibitors in combination drive durable responses by suppressing tumour relapse and inducing development of immune memory, which can also be induced by combination of RAS^G12C^(ON) and PD-1 inhibitors. In contrast, in an immune excluded, “cold”, mouse model of lung cancer, combined RAS^G12C^(ON) and SHP2 inhibition does not cause durable responses, but does sensitise tumours to immune checkpoint blockade, enabling efficient tumour rejection, accompanied by significant TME reorganization, including depletion of immunosuppressive innate immune cells and recruitment and activation of T and NK cells. These preclinical results demonstrate the potential of the combination of RAS^G12C^(ON) inhibitors with SHP2 inhibitors to sensitize anti-PD-1 refractory tumours to immune checkpoint blockade by stimulating anti-tumour immunity as well as by targeting KRAS-driven proliferation in tumour cells.

## Introduction

KRAS oncogenic mutations are very frequent in lung cancer and the leading causes of cancer-related deaths worldwide ^1,2^. KRAS proteins act as signalling hubs where upstream receptor tyrosine kinase (RTK) activation enables guanine exchange factors (GEF) to facilitate the shift of KRAS from the GDP-bound inactive state to the GTP-bound active state ^3^. Glycine to cysteine substitutions at codon 12 (G12C) are the most prevalent KRAS oncogenic mutations in lung cancer ^4^. In contrast to other codon 12 mutations, G12C mutations do not majorly impair the intrinsic GTPase hydrolytic activity of KRAS but do render KRAS^G12C^ insensitive to canonical GTPase-activating proteins (GAP) and thus drive the cycling KRAS^G12C^ to exist predominantly in its active GTP-bound form ^5^. KRAS^G12C^ mutant specific inhibitors such as adagrasib (MRTX849) and sotorasib (AMG 510) have been developed which covalently bind to the mutant cysteine residue of GDP-bound KRAS^G12C^ and block it in its inactive state, preventing it from cycling back to the active GTP-bound state ^6,7^.

The promising initial clinical responses with KRAS^G12C^ inhibitors adagrasib and sotorasib led to their approval for clinical use for advanced KRAS^G12C^ mutant non-small cell lung cancer (NSCLC). However, tumours quite quickly develop resistance to these drugs with median progression free survivals under seven months ^8–10^. Several mechanisms of acquired and adaptive resistance have been suggested which underlines the necessity for developing combination therapies to potentiate the effects of KRAS^G12C^ inhibitors ^11,12^. A number of preclinical studies have shown that KRAS^G12C^ inhibition results in feedback reactivation of upstream RTKs which can result in the activation of wild-type and mutant RAS isoforms. Therefore, inhibition of SHP2, which mediates upstream signalling from RTKs to RAS, is an alternative mechanism to overcome RTK-mediated adaptive resistance ^13–19^. Elevated expression of mutant KRAS in its active form has also been proposed as a mechanism of resistance to KRAS^G12C^ (OFF) inhibitors, thus this could be overcome by using RAS^G12C^(ON) inhibitors ^20^.

Recently, RAS^G12C^ inhibitors have been developed that act as non-degrading molecular glues to create an inactive tri-complex with cyclophilin A (CYPA) and the GTP-bound, active form of KRAS^G12C^. These have been termed RAS^G12C^(ON) inhibitors and include RMC-4998 which is a preclinical tool compound representative of the investigational agent RMC-6291. The binding of these drugs to the active state of KRAS may resolve some limitations of the inactive state selective KRAS^G12C^(OFF) inhibitors ^21^. RMC-6291 is currently undergoing clinical evaluation ^22^ and while duration of response results are awaited, based on experience to date with targeted cancer therapies it is likely that tumours will also develop resistance eventually, meaning that combination strategies will be needed.

The specificity of mutant specific KRAS^G12C^ inhibitors for mutant KRAS in cancer cells without inhibiting wild type KRAS in normal cells has helped to define the role of oncogenic KRAS signalling in modulating anti-tumour immune responses. This includes suppression of interferon signalling, cytokine expression and subsequent recruitment of immunosuppressive myeloid populations and suppression of adaptive T cell-mediated anti-tumour immune responses ^23,24^. Numerous studies have demonstrated that KRAS inhibition reverses immune suppression, causes a profound remodelling of the tumour immune microenvironment (TME) and activates anti-tumour immunity, which provides a window of opportunity for combination with immunotherapies ^24–26^. In contrast with targeted therapies, immunotherapies result in long-term responses in a small fraction of patients ^27^. Moreover, anti-PD-1/anti-PD-L1 therapies are the first line of treatment for advanced NSCLC ^28,29^. Based on that, several clinical trials are exploring combinations of KRAS^G12C^ inhibitors with anti-PD-1/anti-PD-L1 immune checkpoint blockade (ICB). However, using preclinical models, we have recently demonstrated that this combination is only beneficial in immunogenic tumours, characterised by high lymphocyte infiltration and some baseline sensitivity to anti-PD-1, while no combination benefit was observed in immune evasive tumours, which lack lymphocyte infiltration and ICB sensitivity ^24^. Therefore, additional combination therapies that can reshape the TME further may be required to achieve long-term responses using KRAS^G12C^ inhibitors, in particular in patients with immune cold tumours.

SHP2 inhibitors not only suppress the adaptive rewiring of signalling within the cancer cells that diminishes sensitivity to KRAS^G12C^ inhibition, but can also directly affect signalling of non-cancer cells within the TME and enhance anti-tumour immune responses ^15,30–33^. Here, we combine the active state selective RAS^G12C^(ON) specific compound RMC-4998 with the SHP2 inhibitor RMC-4550 and ICB and investigate the effects of these combinations in preclinical models of lung cancer with varying degrees of immunogenicity. We demonstrate that MAPK reactivation in the cancer cells is unavoidable even when using a compound targeting KRAS^G12C^ active state, although this can be suppressed and delayed with the addition of a SHP2 inhibitor. Furthermore, we show that the combination of RMC-4998 and RMC-4550 generates long-term responses in mice with immunogenic lung tumours, including a high proportion of immune mediated tumour eradication. In mice with aggressive immune excluded lung tumours, combination of RMC-4998 and RMC-4550 reshapes the TME, activates adaptive immunity and sensitises tumours to ICB, thus converting these non-inflamed tumours to a more inflamed, ICB-sensitive phenotype. Finally, we provide evidence that suggests direct effects of SHP2 inhibitor on cells of the TME.

## RESULTS

### Combination of RAS^G12C^(ON) and SHP2 inhibition suppresses MAPK reactivation and promotes IFN responses

In order to investigate the tumour cell intrinsic effects of the tri-complex inhibitor targeting RAS^G12C^(ON), RMC-4998, we initially performed cell viability assays using KRAS^G12C^-mutant NSCLC cell lines treated with either RMC-4998 or the KRAS^G12C^(OFF) inhibitor adagrasib (MRTX849). Treatment of both human (CALU1 and NCI-H23) and murine (KPAR^G12C^ and 3LL-ΔNRAS) NSCLC cell lines with either compound led to a concentration-dependent decrease in cell viability, with RMC-4998 displaying increased activity compared to MRTX849 (**Fig. 1a, S1a**). Next, we assessed the effect of both compounds on downstream signalling. MAPK activity was rapidly abrogated with RMC-4998, indicated by a reduction in ERK phosphorylation within the first 15 minutes of incubation, whereas MAPK inhibition with MRTX849 was only evident at later time points (**Fig. S1b, c**). The RAS^G12C^(ON) inhibitor thus acts more rapidly to block KRAS signalling than the KRAS^G12C^(OFF) inhibitor, as expected since it is directly targeting the active form of KRAS. However, at longer time points, targeting the active state of KRAS^G12C^ with RMC-4998 also showed a recovery of ERK phosphorylation at 24-48 hrs that was more comparable to that seen with MRTX849, indicating reactivation of the MAPK pathway and development of adaptive mechanisms (**Fig. 1b**). These observations suggest that even using inhibitors that target the active state of KRAS^G12C^, adaptive mechanisms will eventually develop, emphasizing the necessity for combinatorial strategies.

**Figure 1.**
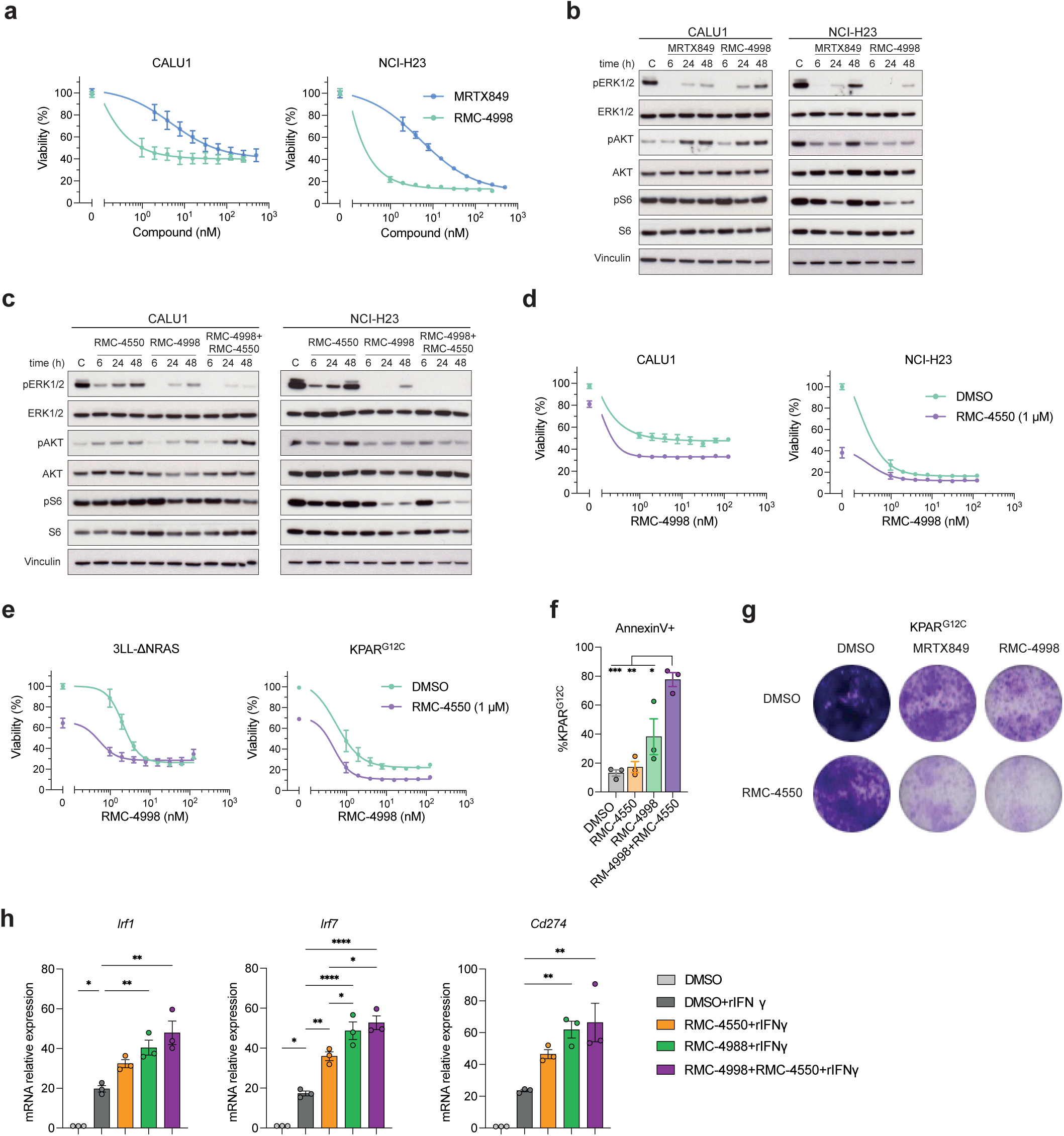
SHP2 inhibitor RMC-4550 prevents adaptive response to the active-state selective RAS^G12C^ inhibitor RMC-4998. (a) Viability of human KRAS-mutant cell lines treated with serial dilutions of RMC-4998 or MRTX849 for 72 hours. Data are mean ± SEM of three independent experiments. (b) Western blot of human KRAS-mutant cell lines treated for 6, 24 or 48 hours with 100 nM MRTX849 or 100 nM RMC-4998. DMSO-treated cells were used as control (C). (c) Western blot of human KRAS-mutant cell lines treated at different time points with 1 µM RMC-4550, 100 nM RMC-4998 or the combination. (d, e) Viability of human (d) or mouse (e) KRAS-mutant cell lines treated for 72 hours with serial dilutions of RMC-4998 in the presence or absence of 1 µM RMC-4550. Data are mean ± SEM of three independent experiments. (f) Percentage of annexin V positive KPAR^G12C^ cells after 72-hour treatment with 1 µM RMC-4550, 100 nM RMC-4998 or the combination. Data are mean ± SD of three independent experiments. Statistics were calculated using one-way ANOVA. (g) Crystal violet staining of KPAR^G12C^ cells treated for 6 days with 200 nM MRTX849, 200 nM RMC-4998, 1 µM RMC-4550 or the combination. (h) qPCR analysis of IFN-induced genes in KPAR^G12C^ cells treated for 24 hours with 1 µM RMC-4550, 100 nM RMC-4998 or the combination, in presence of 100 ng/ml recombinant IFNγ. DMSO treated cells are used as control. Data are mean ± SD of three independent experiments. Statistics were calculated using one-way ANOVA.

Previous studies have demonstrated the role of SHP2 in promoting MAPK pathway reactivation by propagating signalling downstream of elevated upstream RTK activity including inducing cycling of wild type K-, H- and N-RAS isoforms to the GTP-bound activated state ^14^. For this reason, we combined RMC-4550, a SHP2 allosteric inhibitor ^32^, with the RAS^G12C^(ON) inhibitor RMC-4998 to assess if MAPK reactivation could be suppressed. Indeed, in human NSCLC cell lines combination of RMC-4550 with RMC-4998 prevented the rebound of ERK phosphorylation at later time points (24-48h) and resulted in a stronger reduction in cell viability (**Fig. 1c-d**). Moreover, this combination led to enhanced pathway inhibition, shown by decreased expression of downstream MAPK transcript target *Dusp6*, decreased cell viability and increased apoptosis in the mouse cell line KPAR^G12C^ (**Fig. 1e-f, S1d**). These effects extended to longer time points (6 days), where KPAR^G12C^ cultures were replenished with compounds every 48h, validating previous observations that pathway reactivation is not due to reduced compound availability (**Fig. 1g**).

KRAS inhibition can also lead to increased activation of type I and II interferon (IFN) responses in cancer cells, which are crucial for antitumour immune responses ^24^. Consistent with this, treatment with RMC-4998 enhanced IFNγ transcriptional responses measured by the increased expression of IFN target genes (*Irf1*, *Irf7*, *Irf9*, *Cd274*, *B2m*, *H2-d1*) in response to in vitro IFNγ treatment (**Fig. 1h, S1e**). Interestingly, SHP2 inhibition, using RMC-4550, led to similar effects although to a lesser extent, due to a less efficient suppression of the MAPK pathway (**Fig. 1c**).

In summary, these data demonstrate that in KRAS^G12C^ mutant lung cancer cells, targeting together RAS^G12C^(ON) and SHP2 limits the development of adaptive resistance, which results in increased cancer cell apoptosis and tumour cell intrinsic IFN pathway responses.

### RAS^G12C^(ON) and SHP2 inhibition drive immune-dependent cures in an immunogenic model of NSCLC

Given the crucial role of KRAS in modulating antitumour immune responses, we determined the anti-tumour activity of RMC-4998 and RMC-4550 in treating tumours formed by transplantation of KPAR^G12C^ cells, an immunogenic mouse model of NSCLC ^26^. Notable tumour shrinkage and generation of some durable complete regressions (CRs, where no tumour reoccurrence occurs even after treatment withdrawal) of established subcutaneous KPAR^G12C^ tumours in immunocompetent mice were observed with both RMC-4998 and RMC-4550 (**Fig. 2a, b**). Specifically, after only two weeks of treatment, RMC-4998 administration led to generation of 6/8 (75%) CRs whilst RMC-4550 led to generation of 1/7 (14.3%) CRs. Importantly, combination of RMC-4998 and RMC-4550 prevented tumour relapse after treatment withdrawal in all the treated mice, giving 8/8 CRs, confirming the increased activity of the combination (**Fig. 2b**). Mice that had shown complete tumour regression were then rechallenged on the opposite flank with KPAR^G12C^ subcutaneous injections in the absence of further treatment, to assess development of immune memory against the KPAR^G12C^ cancer cells. Indeed, most mice rejected tumour rechallenge and remained tumour free (**Fig. S2a**). These data indicate that both RMC-4550 and RMC-4998 promote induction of antitumour immune responses. Therefore, we assessed the effects of these compounds in immunodeficient subcutaneous KPAR^G12C^ tumour bearing Rag1-/-mice which lack functional T and B lymphocytes and are incapable of inducing adaptive immune responses. Combination of RMC-4550 and RMC-4998 displayed a higher activity than either RMC-4550 or RMC-4998 alone. However, no generation of CRs was observed and all tumours regrew on treatment withdrawal, suggesting that adaptive immunity is essential for generation of long-term CRs (**Fig. 2c, S2b**).

**Figure 2.**
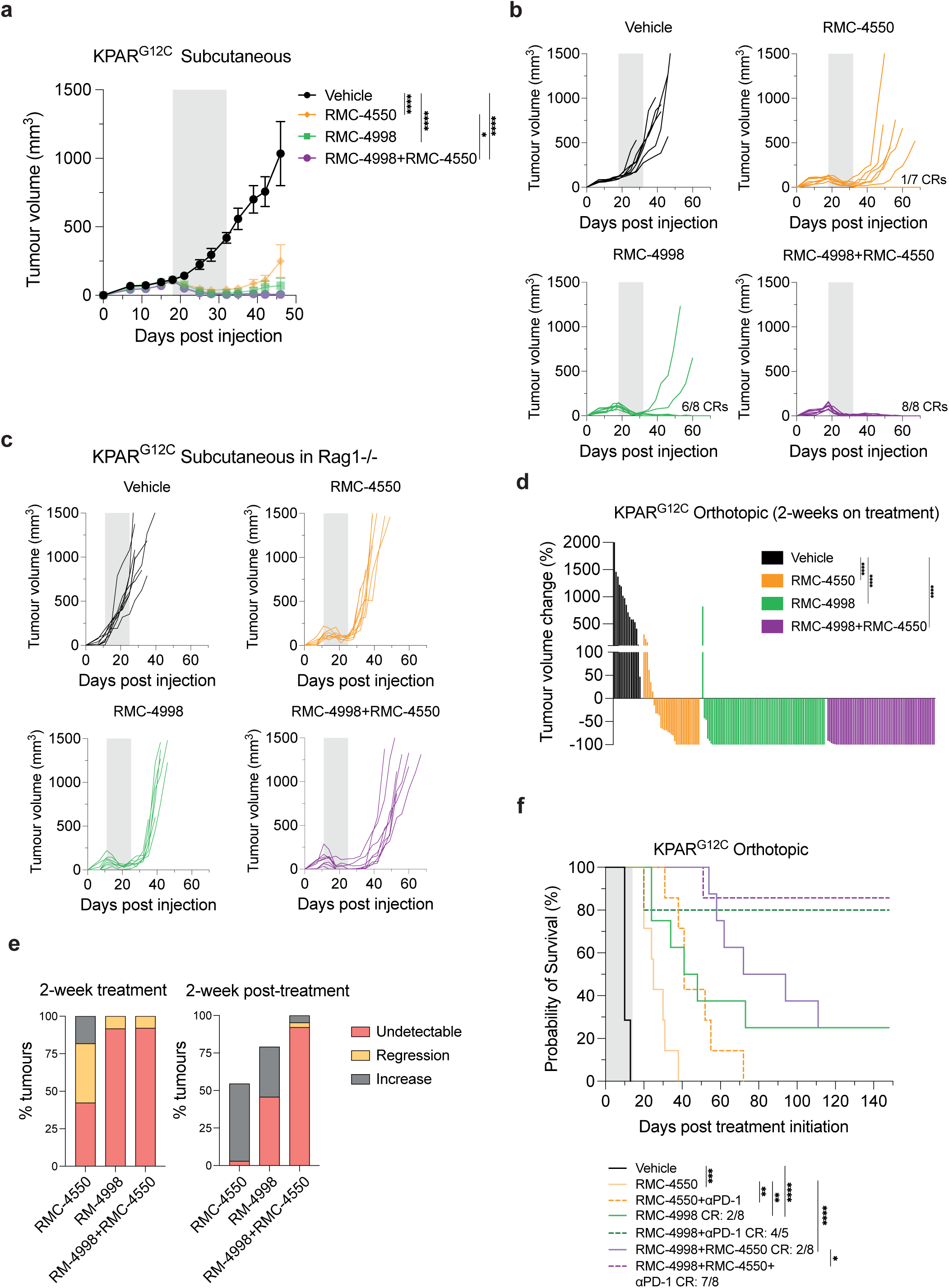
Combination of the RAS^G12C^(ON) inhibitor RMC-4998 with the SHP2 inhibitor RMC-4550 and/or anti-PD-1 in immunogenic KPAR^G12C^ tumours. (a) Tumour growth of KPAR^G12C^ subcutaneous tumours treated daily for 2 weeks with 30 mg/kg RMC-4550 and/or 100 mg/kg RMC-4998. Grey area indicates treatment period. Data are mean tumour volumes ± SEM; n=7-8 mice per group. Analysis was performed using two-way ANOVA. (b) Individual tumour volumes of mice in panel (a). Number of complete regressions (CR) is indicated. Grey area indicates treatment period. (c) Individual tumour growth of KPAR^G12C^ subcutaneous tumours grown in Rag1-/-mice treated daily for 2 weeks with 30 mg/kg RMC-4550 and/or 100 mg/kg RMC-4998. (d) Tumour volume change of orthotopic KPAR^G12C^ tumours after 2 weeks of treatment of mice with 30 mg/kg RMC-4550 and/or 100 mg/kg RMC-4998. Vehicle (n=2), RMC-4550 (n=7), RMC-4998 (n=8), RMC-4998+RMC-4550 (n=8). Each bar represents one tumour. Statistics were calculated using one-way ANOVA. (e) Left panel: Percentage of tumours from panel 2(d) that after two weeks of treatment were undetectable by micro-CT scan, regressing or increasing compared with the initial volume. Right panel: Same tumours were classified based on volume change 2 weeks after treatment withdrawn compared with the tumour volume at the end of the treatment. Missing tumours correspond to mice that died before the 4-week scan. (f) Survival of mice bearing KPAR^G12C^ orthotopic lung tumours treated daily for 2 weeks with 30 mg/kg RMC-4550 and/or 100 mg/kg RMC-4998 in presence or absence of 10 mg/kg anti-PD-1. Grey area indicates treatment period. Number of complete responders (CR) is indicated. n=5-8 mice per group. Analysis was done using log-rank Mantel-Cox test.

Recent studies have highlighted the contrast of adaptive anti-tumour immune responses between subcutaneous and orthotopic lung tumours ^34^. Accordingly, we wanted to extend our findings to the orthotopic setting and investigate the effects of RMC-4550 and RMC-4998 in orthotopic KPAR^G12C^ lung tumours, which have been shown to be partially sensitive to PD-1 blockade ^26^. Similarly to subcutaneous tumours, inhibition of the active form of KRAS^G12C^ achieved better responses than SHP2 inhibition. Micro-CT scanning at the end of the two weeks of treatment revealed that KRAS inhibition with RMC-4998 resulted in 100% regressions in most of the tumours (**Fig. 2d**). Nonetheless, micro-CT scans two weeks after treatment withdrawal revealed that ∼50% of RMC-4998 treated tumours relapsed whereas only ∼10% of RMC-4998 and RMC-4550 combination treated tumours relapsed, demonstrating the importance of this combination in suppressing early tumour relapse (**Fig. 2e**). Treatment of KPAR^G12C^ tumours with RMC-4550 and RMC-4998 for two days resulted in decreased expression of *Dusp6* which indicated suppression of MAPK pathway (**Fig. S2c**), consistent with our in vitro observations (**Fig. S1d**). In parallel, elevated expression of *Cd8a*, *Pdcd1* and *Gzma* and decreased expression of *Arg1* after only two days of treatment suggested infiltration and activation of CD8^+^ T cells and suppression of M2-like myeloid cells, respectively (**Fig. S2c**). Based on these observations and the importance of adaptive immunity for generating long-term responses, we decided to combine RMC-4550 and RMC-4998 with anti-PD-1 immune-checkpoint blockade (ICB) therapy, an established immunotherapy option for NSCLC patients. Notably, addition of anti-PD-1 therapy to RMC-4550 or/and RMC-4998 extended the survival of mice with orthotopic KPAR^G12C^ tumours and resulted in complete regressions in those groups treated with the RAS^G12C^(ON) inhibitor (**Fig. 2f**), even though the period of treatment was only two weeks. Combination of RMC-4998 with anti-PD-1 was the major driver of CRs whilst combination of RMC-4998 and RMC-4550, even though it extended the survival, only generated an equal number of complete responders to RMC-4998 monotherapy.

In summary, these data demonstrate that combination of RAS^G12C^(ON) and SHP2 inhibition can activate adaptive immune responses, suppress tumour relapse, and synergise with anti-PD-1 immunotherapy to generate complete cures at high frequency in an immunogenic mouse model of NSCLC.

### RAS^G12C^(ON) and SHP2 inhibition synergise with ICB to generate cures in a subcutaneous immune-excluded anti-PD-1 resistant model of NSCLC

We next utilised the 3LL-ΔNRAS transplantable mouse model to evaluate responses to RMC-4998 and RMC-4550 and the potential for combination with immunotherapies in an immune excluded, anti-PD-1 resistant lung cancer model. Initially, we assessed the effect of anti-PD-1 and/or anti-CTLA-4 on subcutaneous 3LL-ΔNRAS tumours in immunocompetent mice. Indeed, 3LL-ΔNRAS tumours were completely refractory to anti-PD-1 and/or anti-CTLA-4, with the exception of one responder in the anti-CTLA-4 group (**Fig. 3a, S3a**). Treatment with either RMC-4998 or RMC-4550 resulted in a profound inhibition of tumour growth, which was extended when both compounds were combined. (**Fig. 3b**). In contrast with the results obtained in the KPAR^G12C^ model **(Fig. 2a, b)**, similar responses were generated using either RMC-4998 or RMC-4550. Moreover, treatment with these targeted therapies did not lead to any durable CRs as all mice relapsed while on treatment, including those treated with RMC-4998 and RMC-4550 together (**Fig. 3b, c, S3b**). Of note, addition of anti-PD-1 to the combination of RMC-4998 and RMC-4550 enhanced anti-tumour responses as it led to generation of CRs in 3/8 (37.5%) of mice. In contrast, combination of anti-PD-1 with either RMC-4998 or RMC-4550 alone showed only marginal enhancement of antitumour responses (**Fig. 3d, e**). These data reinforce the previous observations that KRAS^G12C^ inhibition alone does not sensitise immune excluded tumours to anti-PD-1 and underlines the prerequisite for inhibition of both KRAS^G12C^ and SHP2 inhibition for sensitising 3LL-ΔNRAS tumours to anti-PD-1 immunotherapy ^24^. In contrast, both RMC-4998 or RMC-4550 displayed increased anti-tumour activity and induced occasional CRs when combined as single agents with anti-CTLA-4 (**Fig. 3d, e**). However, addition of anti-PD-1 did not much enhance the activity of these doublets (**Fig. 3d, e**), whereas the quadruple combination of targeted and ICB therapies led to tumour eradication in all mice (**Fig. 3c**). These observations are consistent with the hypothesis that the combination of RMC-4998 and RMC-4550 is needed to sensitise tumours to anti-PD-1, and these effects are further enhanced with anti-CTLA-4 treatment. The additional benefit observed with anti-CTLA-4 may reflect depletion of T regulatory cells (Tregs) or attenuation of CTLA-4-mediated inhibition of positive co-stimulation of effector T cells by CD28 ^35^. Finally, the complete responders generated were rechallenged on the opposite flank with 3LL-ΔNRAS subcutaneous injection, without further therapeutic intervention. Importantly, we observed tumour rejection in most mice, which indicated the development of effective immune memory in this extremely immune evasive cancer model **(Fig. S3c, d)**.

**Figure 3.**
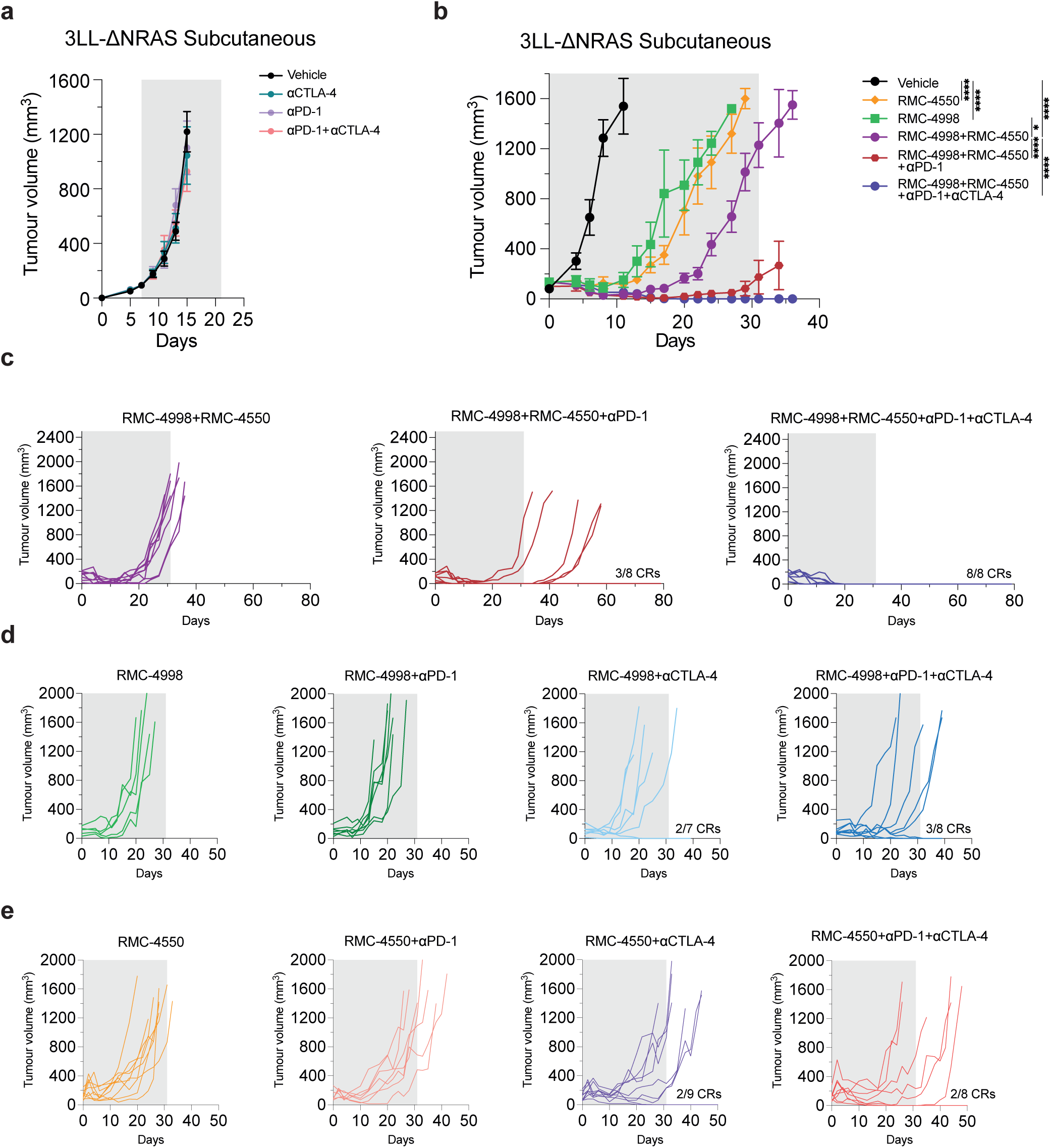
Combination of RAS^G12C^ (ON) inhibitor RMC-4998 with SHP2 inhibitor RMC-4550 sensitises 3LL-Δ NRAS subcutaneous tumours to immunotherapies. (a) Tumour growth of 3LL-ΔNRAS subcutaneous tumours treated with 10 mg/kg anti-PD-1, 5 mg/kg anti-CTLA4 or the combination. Antibodies were administered twice a week for two weeks. Grey area indicates treatment period. Same doses and treatment schedules were administered in the other panels. Data are mean tumour volumes ± SEM; n=6-8 mice per group. Analysis was performed using two-way ANOVA. (b) Tumour growth of 3LL-ΔNRAS subcutaneous tumours treated with 30 mg/kg RMC-4550 and/or 100 mg/kg RMC-4998 in presence or absence of ICB. Grey area indicates treatment period. Data are mean tumour volumes ± SEM; n=7-8 mice per group. Analysis was performed using two-way ANOVA test. (c) Individual tumour volumes of mice in panel (b). Number of complete regressions (CR) is indicated. (d, e) Individual tumour volumes of mice treated with 100 mg/kg RMC-4998 (d) or 30 mg/kg RMC-4550 (e) in presence or absence of ICB. Grey area indicates treatment period. Number of complete regressions (CR) is indicated.

In summary, in the subcutaneous setting, combined RAS^G12C^(ON) and SHP2 inhibition renders immune excluded NSCLC tumours responsive to anti-PD-1 immunotherapy, with further addition of anti-CTLA-4 causing eradication of all tumours.

### RAS^G12C^(ON) and SHP2 inhibition reshape an immune excluded lung cancer TME towards an inflamed phenotype

Previous studies have shown that both KRAS^G12C^(OFF) and SHP2 inhibitors can reshape the immune TME and partially reverse immune suppression ^15,30^. Therefore, we used the orthotopic 3LL-ΔNRAS model to investigate the effects of the RAS^G12C^(ON) inhibitor RMC-4998 and the SHP2 inhibitor RMC-4550 on the lung TME. 3LL-ΔNRAS tumours have been characterised by a predominant infiltration of myeloid cells that suppress immune responses and promote tumour growth. Flow cytometric analysis of established 3LL-ΔNRAS lung tumours validated this along with poor infiltration of lymphocytes (**Fig. 4a**). Immunophenotyping of 3LL-ΔNRAS tumours after 7 days of treatment with RMC-4998 and/or RMC-4550 revealed reduced infiltration of myeloid cells, such as monocytes (CD11b^+^Ly6G^-^Ly6C^+^) and neutrophils (CD11b^+^Ly6G^+^Ly6C^-^; **Fig. 4b**). Different effects between compounds were observed in the CD11b+ tumour-associated macrophages (TAMs), where RMC-4998 increased the infiltration while this population was reduced with the combination (**Fig. 4c**). Additionally, tumours treated with RMC-4998 and/or RMC-4550 had reduced *Arg1* expression, suggesting depletion of Arg1 expressing immunosuppressive myeloid cells (**Fig. 4d**). Interestingly, only the combination of RMC-4998 and RMC-4550 induced elevated expression of *Nos2*, indicating that although either KRAS^G12C^ or SHP2 inhibition can drive depletion of immunosuppressive myeloid-driven programs, only combined inhibition results in the induction of anti-tumour myeloid functions. Considering the heterogeneity of tumour-associated macrophages we aimed to characterise tumour-infiltrating macrophages further. Immunophenotyping of CD11b^+^ TAMs affirmed our observations, as MHCII-low CD206^+^-expressing TAMs were depleted in response to RMC-4998 and/or RMC-4550 treatment (**Fig. 4e**). Moreover, TAMs present in treated tumours expressed higher levels of MHCII with a higher number of them expressing PD-L1, suggesting a functional conversion of TAMs towards antigen-presentation. RNA sequencing (RNA-seq) of treated 3LL-ΔNRAS tumours with either RMC-4998 or RMC-4550 revealed downregulation of myeloid-mediated processes in RMC-4550 treated tumours relative to RMC-4998 (**Fig. S4a**), which is consistent with the differences in infiltration shown in Fig. 4c. Likewise, genes encoding known myeloid-markers were depleted in tumours treated with combined RMC-4998 and RMC-4550 compared to RMC-4998 alone, after 7 days **(Fig. S4b)**. Along with previous reports that SHP2 inhibition can suppress myeloid functions, these data demonstrate additional cancer cell-extrinsic properties of RMC-4550 on myeloid populations ^30,36^.

**Figure 4.**
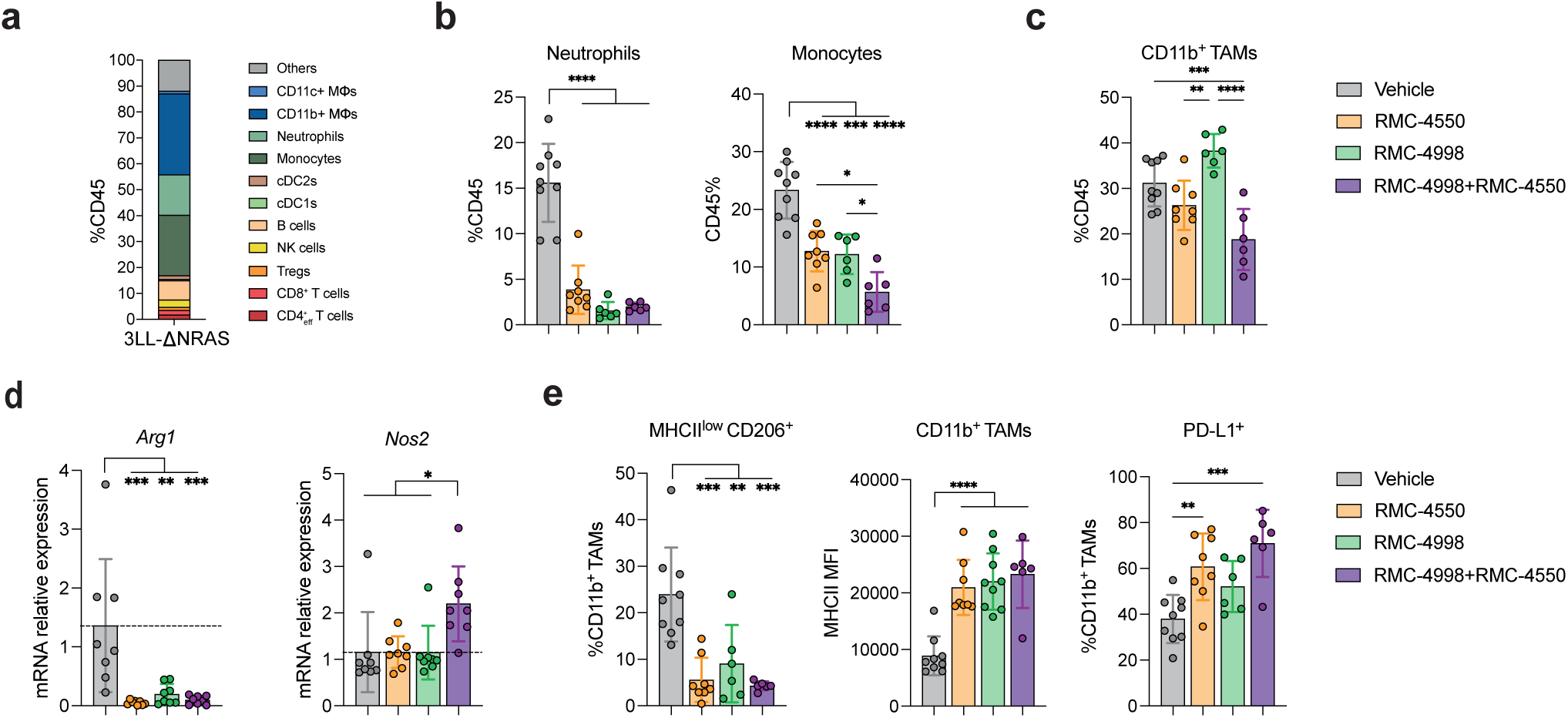
RAS^G12C^ (ON) inhibitor RMC-4998 and SHP2 inhibitor RMC-4550 alter the myeloid cell-landscape in the TME of 3LL-ΔNRAS lung tumours. (a) Flow cytometry immunophenotyping of untreated 3LL-ΔNRAS lung tumours (n=9). (b, c) Frequency of neutrophils, monocytes (b) and CD11b^+^ tumour infiltrating macrophages (c) in 3LL-ΔNRAS lung tumours treated for 7 days with 100 mg/kg RMC-4998, 30 mg/kg RMC-4550 or the combination. Data are mean values ± SD. Each dot represents one mouse. Analysis was performed using one-way ANOVA test. (d) qPCR analysis of Arg1 and Nos2 of 3LL-ΔNRAS lung tumours treated as in (b). Data are mean values ± SD; n = 4 mice per group. Each dot represents one tumour, 2 tumours per mouse. Analysis was performed using one-way ANOVA test. (e) Frequency of MHCII low-expressing and CD206^+^CD11b^+^ tumour infiltrating macrophages (left panel), mean fluorescent intensity of MHCII on CD11b^+^ tumour infiltrating macrophages (middle panel), frequency of PD-L1^+^ CD11b^+^ tumour infiltrating macrophages in 3LL-ΔNRAS lung tumours treated for 7 days with 100 mg/kg RMC-4998, 30 mg/kg RMC-4550 or the combination. Data are mean values ± SD. Each dot represents a mouse. Analysis was performed using one-way ANOVA test.

Next, we examined the impact of RMC-4998 and RMC-4550 treatment on lymphocytes, considering the role of oncogenic KRAS in suppressing the IFN pathway, preventing subsequent antigen presentation and T cell activation, and the critical involvement of SHP2 in modulating functions of lymphocytes ^37,38^. Similar to what has been previously observed with inactive-state KRAS^G12C^ inhibitors, inhibition of active-state KRAS^G12C^ led to increased tumour-infiltrating CD8^+^ T and Treg cells (**Fig. 5a, b**). RMC-4550-treated tumours were characterised by a further elevated presence of T cells compared to tumours treated with RMC-4998, which was also reflected in tumours treated with combined targeted therapy, again pointing towards additional effects of SHP2 inhibition on the TME independent of any effect on cancer cell signalling (**Fig. 5a, b**). All treatments resulted in an increased activation of both CD8^+^ and CD4^+^ T cells and a shift towards an effector-memory phenotype (**Fig. 5c, S5a**). Moreover, tumour-infiltrating T cells in tumours treated with both RMC-4998 and RMC-4550 showed marked activation, proliferation, and expression of potential anti-tumour cytotoxic molecules, such as TNFα, IFNγ and Granzyme B (**Fig. 5d, S5b-d**). Concordant with these data, gene-set enrichment analysis of tumours treated with either RMC-4550 or RMC-4998 revealed activation of IFN responses (**Fig. S5e, f**), while doublet targeted therapy treated tumours showed enrichment of IFN and inflammatory-related pathways compared to either monotherapy (**Fig. 5e**). Likewise, there was an increase of tumour-infiltrating NK (CD49b^+^Nkp46^+^) cells which displayed increased proliferation in response to doublet RMC-4998 and RMC-4550 therapy, even though no differences between RMC-4998 and/or RMC-4550 treated tumours were observed in terms of NK cell numbers, indicating potential increased activity (**Fig. 5f**). Coinciding with this data, tumour-infiltrating NK cells in tumours treated with both RMC-4998 and RMC-4550 showed elevated expression of anti-tumour cytotoxic molecule Granzyme B (**Fig. S5g**). Interestingly, tumour-infiltrating B cells (B220^+^CD19^+^) were uniquely increased in RMC-4550 treated tumours along with an increase of tumour-infiltrating plasma cells (CD138^+^) (**Fig. 5g**), suggesting additional effects of SHP2 inhibition in the B cell populations.

**Figure 5.**
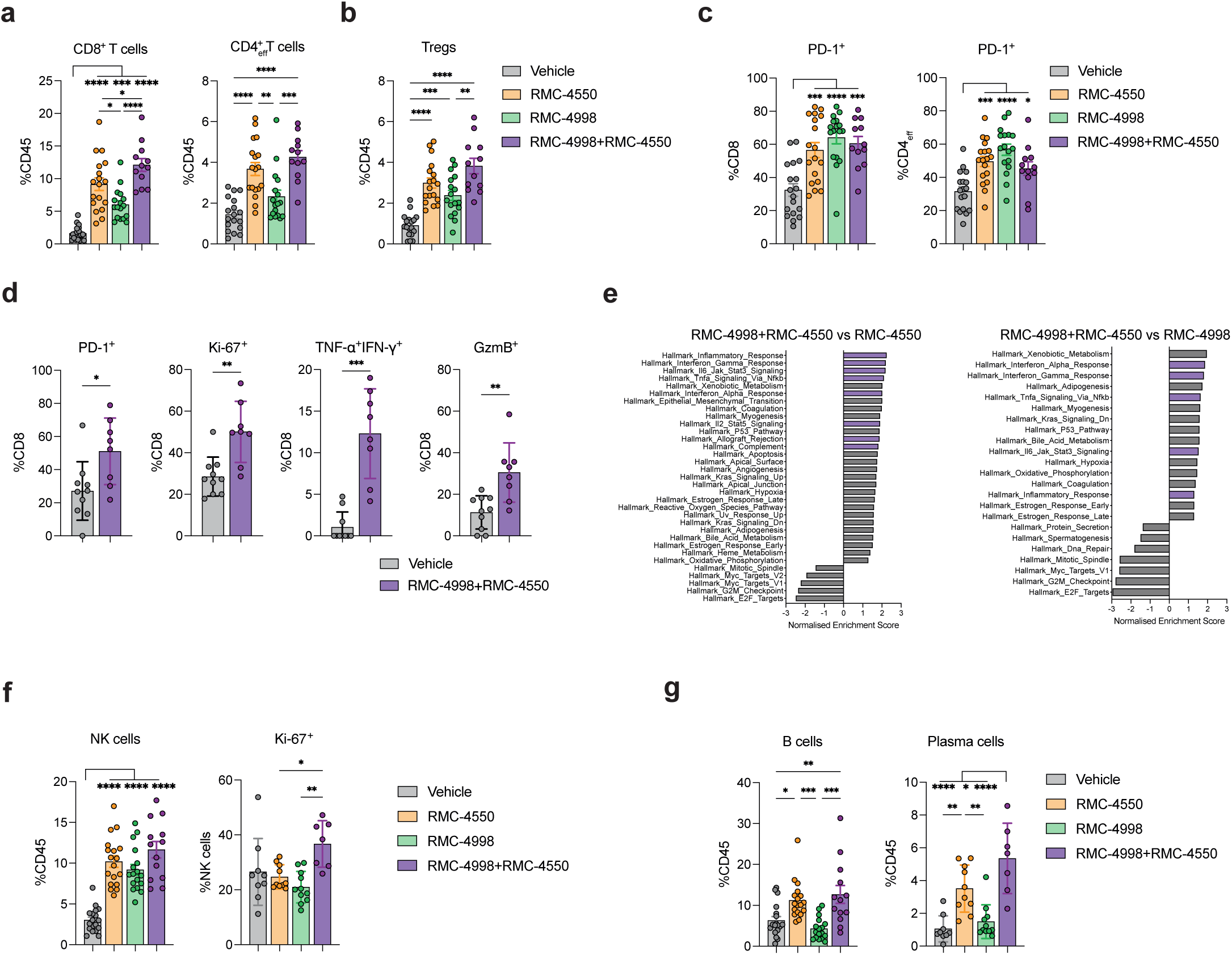
RAS^G12C^(ON) inhibitor RMC-4998 and SHP2 inhibitor RMC-4550 induce pan lymphocytic antitumour response in the TME of 3LL-ΔNRAS lung tumours. (a-c) Frequency of CD8^+^, CD4^+^ effector T (a) and Treg cells (b) and PD-1^+^ CD8^+^, CD4^+^ effector T cells (c) in 3LL-Δ NRAS lung tumours treated for 7 days with 100 mg/kg RMC-4998, 30 mg/kg RMC-4550 or the combination. Each dot represents one mouse. Data ± SEM of two independent experiments. (d) Frequency of PD-1^+^, Ki-67^+^, TNF-α^+^IFN-γ^+^, GzmB^+^ CD8^+^ T cells in 3LL-ΔNRAS lung tumours treated for 7 days with combination of 100 mg/kg RMC-4998 and 30 mg/kg RMC-4550. Each dot represents one mouse. Analysis was performed using two-tailed Student’s t-test. (e) Summary of significantly (FDR < 0.05) down- or upregulated pathways in tumours treated with combined 100 mg/kg RMC-4998 and 30 mg/kg RMC-4550 compared to RMC-4998 or RMC-4550 treated tumours (MSigDB Hallmarks). (f) Frequency of NK and Ki-67^+^ NK cells in 3LL-ΔNRAS lung tumours treated for 7 days with 100 mg/kg RMC-4998, 30 mg/kg RMC-4550 or the combination. Each dot represents one mouse. Data ± SEM of two independent experiments for NK cell frequency. Data ± SD for Ki-67^+^ NK cell frequency. (g) Frequency of B, plasma cells in 3LL-ΔNRAS lung tumours treated for 7 days with 100 mg/kg RMC-4998, 30 mg/kg RMC-4550 or the combination. Each dot represents a mouse. Data ± SEM of two independent experiments for B cell frequency. Data ± SD for plasma cell frequency. Unless otherwise stated, statistics were calculated using one-way ANOVA.

In conclusion, these results underline a mechanistic rationale for combining activestate selective RAS^G12C^ inhibitors, such as RMC-4998, with a second targeted compound, such as the SHP2 inhibitor RMC-4550, that in addition to enhancing the tumour cell intrinsic responses can directly target the TME and potentiate immune responses. Combination treatment with these compounds leads to considerable alteration of the immune TME and results in parallel activation of both innate and adaptive anti-tumour immune responses.

### RAS^G12C^(ON) and SHP2 inhibition synergise with ICB in an orthotopic immune-excluded anti-PD-1 resistant model of NSCLC

As mentioned above, tissue site is an important factor for determining anti-tumour responses. Therefore, we next investigated the therapeutic impact of RMC-4998 and/or RMC-4550 in combination with ICB therapy on the ICB-resistant 3LL-ΔΝRAS model in the orthotopic lung setting. Initially, we confirmed that 3LL-ΔΝRAS lung tumours are refractory to treatment with anti-PD-1, anti-CTLA-4, or the combination of both (**Fig. S6a**). Next, we determined the effect of RMC-4998 and RMC-4550 on tumour growth after 7 days of treatment. Treatment with the SHP2 inhibitor RMC-4550 resulted in a slowing of tumour growth but most of the tumours were still progressing. In contrast, more than half of the tumours treated with the RAS^G12C^(ON) inhibitor RMC-4998 regressed (**Fig. 6a**). Both RMC-4998 and RMC-4550 monotherapies extended survival to comparable levels, which was surprising given the lack of clear regressions induced by RMC-4550 after 7 days of treatment (**Fig. 6a, b**). The long-term effects of RMC-4550 on survival may reflect the non-tumour cell intrinsic effects described earlier, which result in activation of anti-tumour responses and delay of tumour growth. In agreement with the subcutaneous data, combination of RMC-4998 and RMC-4550 enhanced tumour responses, with profound regressions in almost all the tumours analysed and an extension of survival.

**Figure 6.**
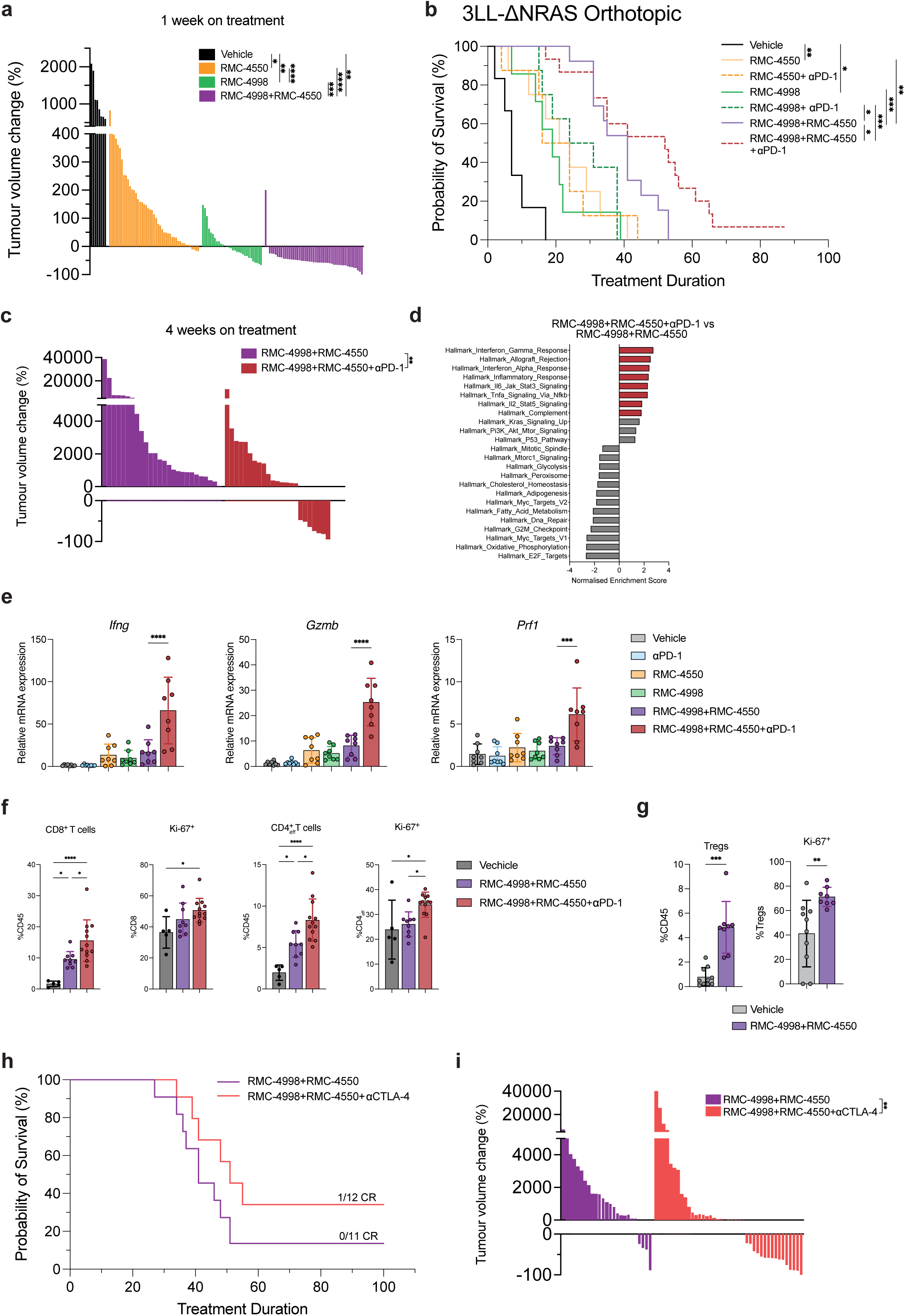
RAS^G12C^(ON) inhibitor RMC-4998 and SHP2 inhibitor RMC-4550 synergise with ICB in an orthotopic immune-excluded anti-PD-1 resistant model of NSCLC. (a) Tumour volume change after 7 days of treatment of 3LL-ΔNRAS tumours with 100 mg/kg RMC-4998 or 30 mg/kg RMC-4550 or the combination. Each bar represents one tumour. Analysis was done using Welch’s one-way ANOVA. (b) Survival of mice bearing 3LL-ΔNRAS orthotopic lung tumours treated daily with 30 mg/kg RMC-4550 or 100 mg/kg RMC-4998 or combination in presence or absence 10 mg/kg anti-PD-1. Vehicle (n=6), RMC-4550 (n=8), RMC-4998 (n=8), RMC-4550+αPD-1 (n=8), RMC-4998+αPD-1 (n=8), RMC-4998+RMC-4550 (n=13), RMC-4998+RMC-4550+α PD-1 (n=15). Analysis was done using log-rank Mantel Cox test. (c) Tumour volume change of mice in panel 6(b) after 4 weeks of treatment of 3LL-ΔNRAS tumours with the combination 100 mg/kg RMC-4998 and 30 mg/kg RMC-4550 in presence or absence of 10 mg/kg anti-PD-1. Each bar represents one tumour. Statistics were calculated using Mann-Whitney test. (d) Summary of significantly (FDR < 0.05) down- or upregulated pathways in tumours treated for 7 days with the triple combination 100 mg/kg RMC-4998 and 30 mg/kg RMC-4550 and 10 mg/kg anti-PD-1 compared tumours treated with combination of 100 mg/kg RMC-4998 and 30 mg/kg RMC-4550 (MSigDB Hallmarks). (e) qPCR analysis of immune genes of 3LL-ΔNRAS orthotopic lung tumours treated for 7 days with 10 mg/kg anti-PD-1, 30 mg/kg RMC-4550, 100 mg/kg RMC-4998, the dual combination of RMC-4998 + RMC-4550 or the triple combination RMC-4998 + RMC-4550 + anti-PD-1. Each dot represents one tumour, 2 tumours per mouse. Statistics were calculated using one-way ANOVA. (f) Frequency of CD8^+^ and CD4^+^ T cells and frequency of Ki-67^+^ CD8^+^ and CD4^+^ T cells in 3LL-ΔNRAS tumours after 2 weeks of treatment with the combination of 100 mg/kg RMC-4998 and 30 mg/kg RMC-4550 in presence or absence of 10 mg/kg anti-PD-1. Each dot represents one mouse. (g) Frequency of Treg cells and Ki-67^+^ Treg cells in 3LL-ΔNRAS lung tumours treated for 7 days with combination of 100 mg/kg RMC-4998 and 30 mg/kg RMC-4550. Each dot represents one mouse. Analysis was performed using two-tailed Student’s t-test. (h) Survival of mice bearing 3LL-ΔNRAS orthotopic lung tumours treated with the combination 100 mg/kg RMC-4998 and 30 mg/kg RMC-4550 in presence or absence of 10 mg/kg anti-CTLA-4 (n=11-12). CRs, where no tumour was detected by microCT scan one month after treatment withdrawal. (i) Tumour volume change after 4 weeks of treatment of 3LL-ΔNRAS tumours with the combination 100 mg/kg RMC-4998 and 30 mg/kg RMC-4550 in presence or absence of 10 mg/kg anti-CTLA-4. Each bar represents one tumour. Statistics were calculated using Mann-Whitney test.

Although treatment of tumours with either RMC-4998 or RMC-4550 monotherapy extended survival and enhanced the inflamed phenotype of the TME, addition of anti-PD-1 to either alone did not improve survival responses. In contrast, the targeted therapy doublet sensitised tumours to anti-PD-1, leading to significantly prolonged survival, consistent with our previous observations in subcutaneous tumours (**Fig. 6b, 3c-e**). Micro-CT scans after 4 weeks on treatment revealed that combination of anti-PD-1 with doublet targeted therapy suppressed tumours more efficiently and prevented them from growing back (**Fig. 6c**). Moreover, RNA-seq of 7-day treated tumours indicated an enrichment of IFN and inflammatory-related pathways in tumours treated with triple combination of RMC-4998, RMC-4550 and anti-PD-1 compared to dual targeted therapy (**Fig. 6d**). In agreement with these data, qPCR of treated tumours demonstrated a profound increased expression of genes involved in IFN pathway and cytotoxic response (*Ifng*, *Gzmb*, *Prf1*; **Fig. 6e**). Additionally, tumours treated with triple combination for two weeks were characterised by an elevated infiltration and persistent proliferation of CD8^+^ and CD4^+^ T cells (**Fig. 6f**). These data support the idea that combination of RMC-4998 and RMC-4550 can sensitise immune excluded tumours to anti-PD-1 immunotherapy through enhanced IFN pathway induction and T cell persistence.

However, the triple combination still failed to generate complete responses in the orthotopic lung setting, in contrast to the subcutaneous tumours (**Fig. 3c**). To extend our findings, we decided to assess if the combination of RMC-4998 and RMC-4550 could sensitise tumours to other immunotherapies and generate complete responses. In 3LL- ΔΝRAS lung tumours, both RMC-4998 and/or RMC-4550 treatments induced infiltration and activation of Treg cells (**Fig. 5b, 6g**). It has been previously shown that Tregs can dampen anti-tumoural responses, prompting us to investigate treatments that may impact on Treg cells. We employed an anti-CTLA-4 depleting antibody to deplete T regs and attenuate CTLA-4 inhibition of positive co-stimulation by CD28. In combination with RMC-4998 and RMC-4550, anti-CTLA-4 further suppressed tumour relapse and extended survival compared with dual targeted therapy while also generating one tumour-free mouse (**Fig. 6h, i**).

These data demonstrate the potential of combining RMC-4998 and RMC-4550 to enhance tumour regressions and prolong survival in an immune excluded, ICB-resistant NSCLC model. In parallel, this combination can be combined further with immunotherapies to generate long term responses through enhancement of IFN responses and anti-tumour immunity.

## Discussion

The short-lived responses observed in patients treated with adagrasib and sotorasib, which target the inactive conformation of KRAS^G12C^, highlight the need for development of therapies that can prevent resistance emergence and prolong responses ^8,9^. In this study we used RMC-4998, a compound that targets the active, GTP-bound form of KRAS^G12C^, and demonstrates a more rapid and potent activity compared to adagrasib. This can be attributed to the limitations of adagrasib, and other inhibitors targeting GDP-bound KRAS^G12C^, which depend on the intrinsic GTPase hydrolysis rate of KRAS^G12C^ to convert KRAS to the inactive state. However, while the use of active state RAS^G12C^ inhibitors, as exemplified with the preclinical tool compound RMC-4998, should prevent certain resistance mechanisms, such as an increase of the active KRAS^G12C^ pool ^20^, the present preclinical data demonstrate that adaptive resistance and MAPK reactivation, potentially caused by the feedback activation of wild-type RAS isoforms, can still be observed ^14^. Here, we show that simultaneously inhibiting SHP2 using RMC-4550 suppressed MAPK reactivation, enhanced induction of apoptosis and maintained IFN pathway activation. Given the role of oncogenic KRAS in driving immune suppression, prevention of MAPK reactivation is crucial to block cancer cell-intrinsic adaptive resistance and the maintenance of an immunosuppressive TME and the potential for cross-resistance ^39^.

Recent studies have demonstrated the paradoxical role of the IFN pathway in cancer. We have previously shown that activation of cancer cell-intrinsic IFN response is crucial for induction of adaptive cytotoxic anti-tumour immune programmes and long-term responses induced by KRAS^G12C^ inhibition, especially in combination with anti-PD-1 therapies ^24^. Here, using an immunogenic lung cancer model, we also demonstrate that adaptive immunity is essential for prevention of relapse and generation of immune memory upon inhibition of KRAS and/or SHP2, highlighting the importance of adaptive immunity in generating longterm anti-tumour responses. In fact, while addition of a SHP2 inhibitor along with RAS^G12C^(ON) inhibition provides a survival benefit, it is the combination of RAS^G12C^(ON) inhibition with anti-PD-1 blockade that is the major driver for tumour eradication in preclinical models with a degree of intrinsic sensitivity to ICB, confirming the critical role of adaptive immunity in the generation of durable responses. However, only a fraction of NSCLC patients benefit from ICB. Moreover, in preclinical models, tumours resistant to ICB do not gain any advantage from the combination of adagrasib and anti-PD-1, even though KRAS^G12C^ inhibition can partially reverse immune suppression mechanisms in these models^24^. Similarly, we do not observe an extension of survival when combining the active-state RAS^G12C^ inhibitor RMC-4998 with anti-PD-1 in 3LL-ΔNRAS tumours, an immune excluded model representative of lung tumours that are intrinsically resistant to ICB. In contrast, in this preclinical model the combination of KRAS^G12C^ and SHP2 inhibitors significantly extends survival, suggesting that patients with immune evasive tumours may obtain greater benefits from this combination. However, indicative of the immunosuppressive nature of this model and in contrast to the immunogenic KPAR^G12C^ model, combined inhibition of both KRAS^G12C^ and SHP2 did not lead to tumour eradication, just delayed progression, suggestive of inadequate engagement of adaptive immunity. Nevertheless, when anti-PD-1 is added in this context, it leads to tumour eradication in some mice and development of immune memory, indicating that targeting both KRAS^G12C^ and SHP2 can sensitise non-immunogenic tumours to anti-PD-1 therapies and achieve long-term responses. In terms of response to immune checkpoint blockade, combination treatment with RAS^G12C^(ON) and SHP2 inhibitors appears to convert immune excluded, “cold” tumours to an immune inflamed, “hot”, state.

Given the pleiotropic role of both oncogenic KRAS and SHP2 in modulating cancer-cell intrinsic mechanisms, including indirect regulation of the TME, and the cancer-cell extrinsic functions of SHP2, there are several potential mechanisms by which combined inhibition of KRAS^G12C^ and SHP2 could sensitise immune evasive tumours to anti-PD-1 therapies. In cancer cells, combination of RAS^G12C^(ON) and SHP2 inhibitors prevents MAPK reactivation, which extends the reversion of immune evasive mechanisms driven by oncogenic KRAS, such as secretion of immune evasive cytokines and inhibition of IFN responses. Moreover, the combination results in elevated cancer cell death which could lead to increased uptake of immunogenic antigens due to phagocytosis by antigen-presenting cells ^40,41^. The increase of antigen presentation along with persistent and increased IFN signalling could prime T cell responses and thus increase the dependency on the anti-PD-1/PD-L1 axis ^42^. In parallel, enhanced tumour regressions and prevention of relapse could expand the T-cell reinvigoration window ^43^. Alternatively, cancer-cell independent effects of SHP2 inhibition on the TME could also contribute to the sensitivity to anti-PD-1. Consistent with our data, it has been shown that SHP2 inhibition depletes immunosuppressive tumour-associated macrophages, which consequently enables recruitment of macrophages with the potential of promoting antitumour immunity ^30^. Recent studies have further characterised the effects of SHP2 inhibition in macrophages which can lead to antitumour immune responses. These include induction of CXCL9 expression, a strong prognostic biomarker in several tumours, and differentiation of bone marrow-derived myeloid cells, both of which can induce CD8^+^ T cells recruitment and activation ^36,44,45^. SHP2 cancer-cell extrinsic effects are not restricted to myeloid cells. In fact, we have observed increased T cell tumour infiltration upon SHP2 inhibition compared to KRAS^G12C^ inhibition. This indicates a direct role of SHP2 in T cells in terms of proliferation and/or recruitment, in agreement with previous reports ^46^. Additionally, we observe increased NK cell expansion and cytotoxicity in tumours treated with doublet therapy. Niogret et al., have shown that SHP2 is indispensable for IL-15R-mediated expansion and activation of NK cells ^37^. Finally, in this study, we present evidence that SHP2 inhibition can promote increased frequency of tumour-infiltrating B cells and plasma cells. B cells have been described to have a crucial role in modulating anti-tumour immunity, including antigen presentation and antibody secretion against tumour-derived retroviral elements ^47,48^.

Anti-PD-(L)1 blocking antibodies, either as single therapy or in combination with chemotherapy, constitute the first line treatment for the majority of patients with KRAS mutant NSCLC ^29^. However, recent studies have also suggested the potential benefits of other immunotherapies, including anti-CTLA-4 blocking antibodies ^49,50^. In this study, we demonstrate that the combination of RAS^G12C^(ON) and SHP2 inhibition also sensitises immune evasive tumours to anti-CTLA-4 treatment, leading to durable responses.

Interestingly, anti-CTLA-4 treatment outperforms anti-PD-1 blockade in subcutaneous tumours. Based on previous observations, we speculate that this differential effect may, in part, be attributed to the capacity of anti-CTLA-4 to deplete FoxP3^+^ T regulatory (Treg) cells ^26,35^. It is well established that Treg accumulation within tumours suppresses anti-tumour immunity through several mechanisms. However, recent studies have identified novel properties of anti-CTLA-4 treatment in activating anti-tumour immune responses, such as Fcγ receptor engagement-mediated myeloid activation ^51^. Therefore, we cannot exclude additional TME influences by anti-CTLA-4 treatment. Given the numerous changes in immune cell populations within the TME induced by the combined inhibition of KRAS and SHP2, this provides rationale for combination with other immunotherapies.

Preliminary clinical activity data have been reported for the investigational agent RMC-6291, an active-state RAS^G12C^ inhibitor, including evidence of clinical activity in patients with advanced KRAS^G12C^ mutant NSCLC that had previously progressed on inactive-state KRAS^G12C^ inhibitors ^22^. It is still too early to know how the duration of the responses to this new generation of RAS^G12C^(ON) inhibitors will compare to those of the first generation of KRAS^G12C^(OFF) inhibitors. However, based on previous experience with targeted therapies, it is likely that resistance will eventually develop and that combinations will be needed. Combinations that maintain or even enhance the positive effects that the mutant specific KRAS inhibitors have in the TME are of particular interest. Here we show that for immune evasive tumours, this can be achieved with additional targeting of SHP2 and that it is the combination of both the tumour cell-intrinsic and cell-extrinsic effects of SHP2 inhibition that sensitises immune evasive tumours to ICB and generates durable responses. It is important to consider the potential toxicities of these combinations. Initial reports from clinical trials of sotorasib combined with anti-PD-1 checkpoint blockade have indicated high incidence of toxicities, in particular immune related hepatoxicity ^52,53^. The mechanism(s) underlying these toxicities is still unclear and requires further investigation. However, emergent data with other KRAS^G12C^(OFF) inhibitors suggests that the effects are likely to reflect compound specific, off target effects, thus allowing hope that combinations involving future generations of KRAS inhibitors could potentially be less toxic. In parallel, studies that analyse the potential of different treatment schedules to maximise the effects of the different therapies reducing the toxicities should be performed. Our studies of combination therapies targeting RAS^G12C^(ON), SHP2 and immune checkpoints in preclinical models support clinical evaluation to assess their potential for improving clinical outcomes in lung cancer, providing the challenges of avoiding adverse toxicities can be addressed.

## METHODS

### Sex as a biological variable

The KPAR^G12C^ cell line was generated from a female mouse whereas the 3LL-ΔNRAS cell line was derived from a male mouse. Our study examined antitumour immune responses generated by targeted therapy and to avoid introducing error due to induction of immune responses in female mice against genes found in the Y chromosome, we used mice with the same sex as the sex of the mouse that the cell lines were derived from, i.e. male mice for transplantation of 3LL-ΔNRAS cells and female mice for transplantation of KPAR^G12C^ cells. Our study therefore examined both male and female animals, but in different se6ngs, so it is unknown whether the specific findings are relevant for the opposite sex.

### In vivo tumour studies

All studies were performed under a UK Home Office–approved project license and in accordance with institutional welfare guidelines. All transplantation animal experiments were carried out using 8-10-week C57BL/6J mice.

For subcutaneous tumour injections, 150,000 KPAR^G12C^ or 400,000 3LL-ΔNRAS cells were mixed 1:1 with Geltrex LDEV-Free Reduced Growth Factor matrix (Thermo Fisher Scientific) and injected subcutaneously into one flank. Tumour growth was followed two or three times a week by calliper measurements and volume was calculated using the formula 0.5 x (length x width^2^). Mice were euthanised when average tumour diameter exceeded 1.5 cm. Diameter was measured using callipers. For rechallenge experiments, mice with undetectable tumours at least 30 days after the treatment withdrawn were injected subcutaneously with the same number of cells in the opposite flank. Naïve mice of similar age were injected as control.

For orthotopic tumours, 150,000 KPAR^G12C^ or 10^6^ 3LL-ΔNRAS cells were injected in 100 μl of phosphate-buffered saline (PBS) in the tail vein. Tumour volume was measured by micro-CT analysis. Mice were anaesthetised by inhalation of isoflurane and scanned using the Quantum GX2 micro-CT imaging system (Perkin Elmer). Serial lung images were reconstructed and tumour volumes analysed using Analyse (AnalyzeDirect) as previously described ^54^. Mice were weighed and monitored regularly and were euthanized when the humane endpoint of 15% weight loss from baseline was reached or any sign of distress was observed (i.e. hunched, piloerection, difficulty of breathing). In addition, if a mouse was observed to have a tumour burden in excess of 70% of lung volume when assessed by micro-CT scanning, they were deemed at risk of rapid deterioration in health and euthanised immediately.

Mice were randomized into groups and treatments were initiated once tumours reached an average volume of 50-150 mm^3^ for subcutaneous studies or were detectable by micro-CT for orthotopic experiments. Mice were treated daily via oral gavage with 100 mg/kg RMC-4998 and/or 30 mg/kg RMC-4550. RMC-4998 and RMC-4550 were provided by Revolution Medicines, Inc. under a former collaboration agreement with Sanofi. Compounds were prepared using a formulation made of 10% DMSO / 20% PEG 400 / 10% Kolliphor HS15 in 50 mM sodium citrate buffer pH 4. For ICB treatments, mice were administered with 10 mg/kg anti-PD-1 (clone RMP1-14, cat. BE0146, BioXcell) and/or 5 mg/kg anti-CTLA-4 (clone 9H10, cat. BE0131, BioXcell). Antibodies were dissolved in PBS and administered via intraperitoneal injection (4 μl/g) twice weekly for two weeks. For 3LL-1′NRAS orthotopic experiments, mice were treated with 10mg/kg anti-PD-1 (clone RMP1-14, cat. mpd1-mab15, InvivoGen) or 10 mg/kg anti-CTLA-4 (clone 9D9, cat. mctla4-mab10, InvivoGen). Antibodies were administered twice weekly for a maximum of four weeks.

### Cell lines and treatments

NCI-H23 and Calu-1 were obtained from the Francis Crick Institute. 3LL-1′NRAS cells were generated as previously described ^55^. KPAR1.3 G12C cells (herein KPAR^G12C^) were generated as previously described ^26^. NCI-H23 cells were grown in RPMI and the rest of the cells in DMEM. Medium was supplemented with 10% fetal calf serum, 4 mM L-glutamine, penicillin (100 U/ml) and streptomycin (100 mg/ml). Cell lines were routinely tested for mycoplasma and were authenticated by short-tandem repeat (STR) DNA profiling by the Francis Crick Institute Cell Services facility.

Cells were plated at an appropriate density and left to grow for at least 24 hours before treatment. RMC-4998 and RMC-4550 were provided by Revolution Medicines under a sponsored research agreement. MRTX849 was obtained from MedChemExpress. Recombinant mouse IFNγ was obtained from Biolegend.

### Cell viability and apoptosis assays

For viability assays, cells were grown in 96-well plates and inhibitors were added 24 hours later. After 72 hours, 5 μl of CellTiter-Blue (Promega) was added and cells were incubated for 90 minutes at 37°C before measuring fluorescence using an EnVision plate reader (PerkinElmer). For longer-term proliferation assays, cells were plated in 24-well plates and treated for 6 days. Compounds were replaced after 3 days of treatment. At the end of treatment, cells were fixed and stained using 0.2% crystal violet in 2% ethanol.

Percentage of apoptotic cells was measured using flow cytometry. Cells were plated in 6-well plates and treated for 72 hours. At the end of the treatment, harvested cells were resuspended in Annexin V binding buffer and stained with FITC Annexin V (BD Biosciences) and DAPI.

### Western Blotting

Cells were plated in 6-well plates and treated 24 hours later. At the end of the treatment, cells were lysed using 10X Cell Lysis Buffer (Cell Signaling) supplemented with Complete Mini protease inhibitor cocktail and PhosSTOP phosphatase inhibitors (Roche). After quantification using a BCA protein assay kit (Pierce), proteins (15-20 μg) were separated on 4-12% NuPAGE Bis-Tris gels (Life Technologies) followed by transfer to PVDF membranes. Bound primary antibodies were incubated with HRP-conjugated secondary antibodies (Amersham) and detected using chemiluminescence (Luminata HRP substrate, Millipore). List of antibodies used is listed in **Supplementary Information 1**.

## Quantitative RT-PCR

RNA was extracted from cell lines or frozen lung tumours using the RNeasy Mini Kit (Qiagen) following the manufacturer’s instructions. For in vivo tumour samples, tumours individually isolated from lungs were lysed and homogenized using RNase-free disposable pellet pestles (Kimble Chase) followed by QIAshredder columns (Qiagen). cDNA was generated using the Maxima First Strand cDNA Synthesis Kit (Thermo Fisher Scientific) and qPCR performed using Fast SYBR Green Master Mix (Applied Biosystems). List of primers used is detailed in **Table 1**. Gene expression changes relative to the housekeeping genes were calculated using the ΔΔCT method.

**Table 1.**
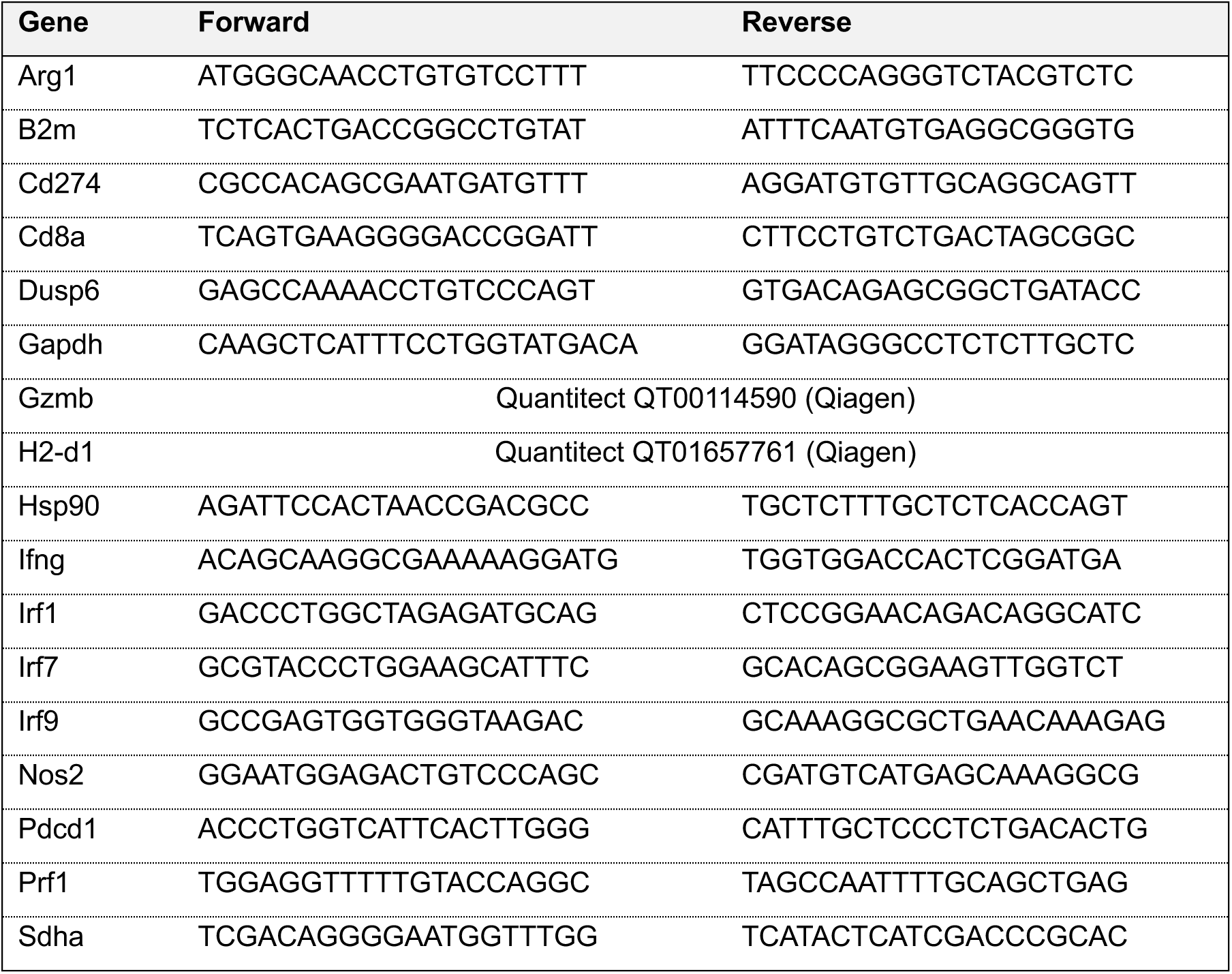
List of qPCR primers.

### RNA sequencing

RNA was extracted as indicated above. RNA quality was measured using the TapeStation 4000 (Agilent). Libraries were prepared using the NEBNext Ultra II Directional PolyA mRNA (New England Biolabs) according to manufacturer’s instructions, with an input of 150 ng and 10 PCR cycles for library amplification. Library quality was measured using the TapeStation 4200 (Agilent). Samples were sequenced to a depth of ∼25M paired-end 100 bp reads in an Illumina NovaSeq 6000 system.

For data analysis, sequencing reads were pre-processed and mapped to the mouse GRCm38 genome using the nfcore/rnaseq pipeline with STAR and RSEM. The R package DESeq2 was used to perform differential analysis between treatment groups. Gene set enrichment analysis was carried out with the R package fgsea using a ranked list of Wald statistic values calculated by DESeq2 for each comparison. Genes were taken from the Hallmark set available at MSigDB using the R package msigdbr.

### Flow cytometry

Mice were euthanised using schedule 1 methods, lung tumours were dissociated and all tumours from one lung were pooled together. Tumours were finely cut into small pieces and digested with collagenase (1 mg/ml; Thermo Fisher Scientific) and DNase I (50 U/ml; Life Technologies) in HBSS for 45 min at 37°C. Samples were filtered through 70 μm strainers (Falcon) and red blood cells were lysed using ACK buffer (Life Technologies). After washes in PBS, cells were stained with fixable viability dye eFluor870 (BD Horizon) for 30 minutes and blocked with CD16/32 antibody (BioLegend) for 10 minutes. Samples were washed three times in FACS buffer (2 mM EDTA and 0.5% bovine serum albumin in PBS, pH 7.2) and stained using fluorescently labelled antibody mixes for surface markers. After staining, samples were fixed in Fix/lyse solution (eBioscience). If intracellular staining was performed, cells were instead fixed and permeabilized with Foxp3 / Transcription Factor Fixation/Permeabilization kit (Invitrogen) according to manufacturer’s instructions before staining with intracellular antibodies. Single stain controls with spleen or OneComp eBeads (Invitrogen) were also performed. List of antibodies used is detailed in **Supplementary Information 1**. Samples were resuspended in FACS buffer and analysed using a FACSymphony cytometer (BD). Data was analysed using FlowJo software, detailed in **Supplementary Figure 7**. For intracellular staining of NK and T cells, cell suspensions were incubated in RPMI supplemented with 1:1000 BD GolgiPlug (BD Biosciences), 1 μg/ml ionomycin, 50 ng/ml PMA (all Sigma) for 1 and 4 hours, respectively, and then staining was performed using the Foxp3 / Transcription Factor Fixation/Permeabilization kit (Invitrogen).

### Statistical analysis

Data was analysed using Prism 8 (GraphPad Software) using normality distribution tests, Mantel-Cox, two-tailed Student’s t-test, one-way, two-way or Welch’s ANOVA, as indicated. Significance was determined at p < 0.05 (*p < 0.05, **p < 0.01, ***p < 0.001, ****p < 0.0001).

## ACKNOWLEDGEMENTS

We thank the science technology platforms at the Francis Crick Institute including Biological Resources, Advanced Sequencing, Scientific Computing, Bioinformatics and Biostatistics, Flow Cytometry, Experimental Histopathology, and Cell Services. We thank the members of the Oncogene Biology laboratory for their discussions and critical reading of the manuscript.

## Author contributions

P.A., M.M-A. and J.D. designed the study, interpreted the results and wrote the manuscript. P.A., S.R., A.dC., M.T., J.B., E.M. and S.dC. performed the biochemical experiments. C.M. assisted with *in vivo* studies. R.G. performed bioinformatic analysis. C.B., E.Q. and J.A.M.S. contributed with interpretation and resources. All authors contributed to manuscript revision and review.

---

**Supplementary Figure 1.**
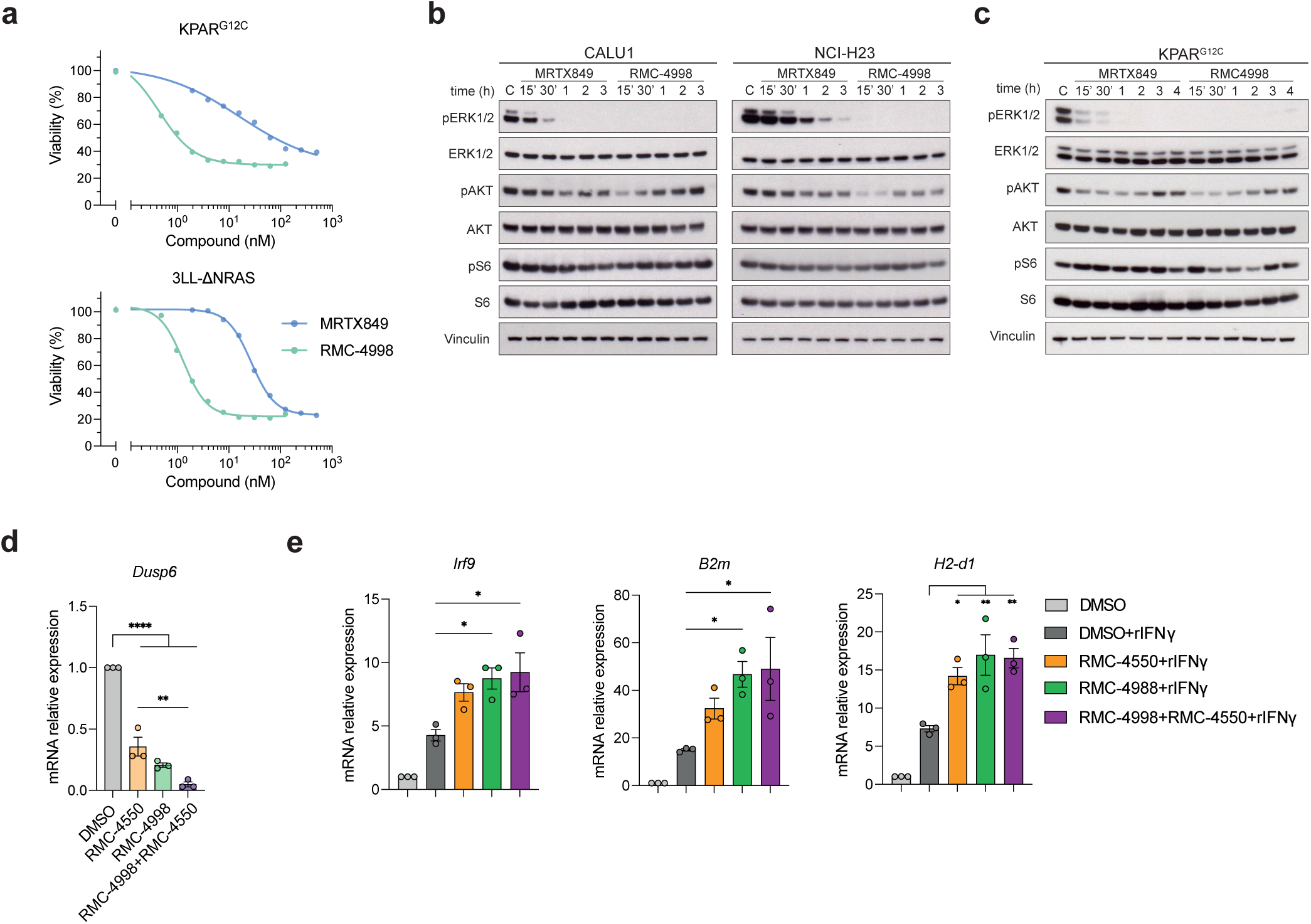
SHP2 inhibitor RMC-4550 prevents adaptive response to the active-state selective RAS^G12C^ inhibitor RMC-4998. (a) Viability of mouse KRAS-mutant cell lines treated with serial dilutions of RMC-4998 or MRTX849 for 72 hours. Data are mean ± SEM of three independent experiments. (b) Western blot of human KRAS-mutant cell lines treated for 15 min, 30 min, 1, 2 or 3 hours with 100 nM MRTX849 or 100 nM RMC-4998. DMSO-treated cells were used a control (C). (c) Western blot of mouse KRAS-mutant cell line treated for 15 min, 30 min, 1, 2 or 4 hours with 100 nM MRTX849 or 100 nM RMC-4998. (d) qPCR analysis of Dusp6 in KPAR^G12C^ cells treated for 24 hours with 1 µM RMC-4550, 100 nM RMC-4998 or the combination. (e) qPCR analysis of IFN-induced genes in KPAR^G12C^ cells treated for 24 hours with 1 µM RMC-4550, 100 nM RMC-4998 or the combination, in presence of 100 ng/ml IFNγ. DMSO treated cells are used as control. Data are mean ± SD of three independent experiments. Statistics were calculated using one-way ANOVA.

**Supplementary Figure 2.**
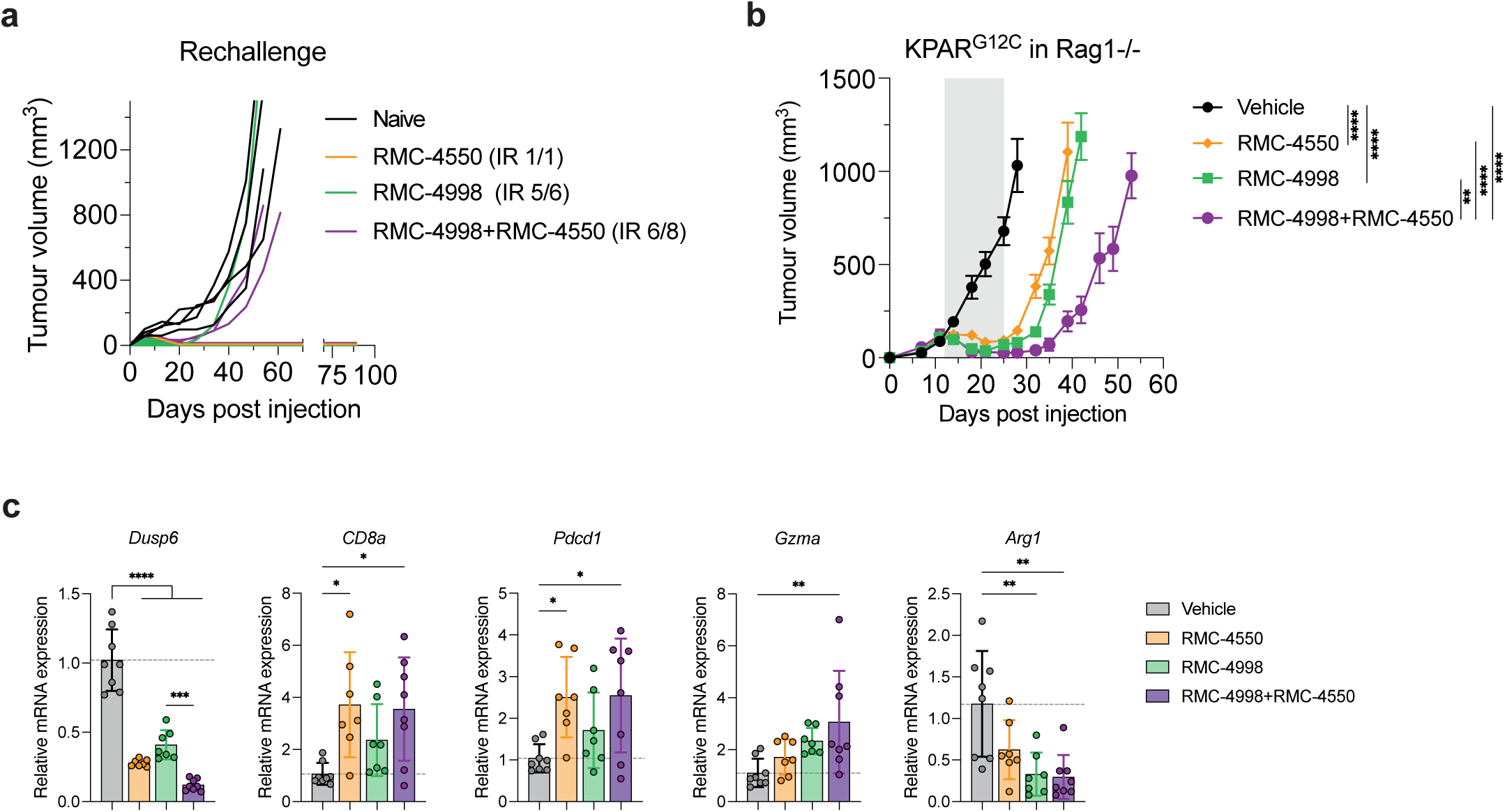
Combination of the RAS^G12C^(ON) inhibitor RMC-4998 with the SHP2 inhibitor RMC-4550 and/or anti-PD-1 in immunogenic KPAR^G12C^ tumours. (a) Mice in Fig. 2a that rejected the primary tumour were rechallenged on the opposite flank and tumour volume was measured. Number of mice that achieved immune rejections (IR) is indicated. Naïve mice of similar age were used as control. Legends indicate the treatment that the primary tumour received. (b) Tumour growth of KPAR^G12C^ subcutaneous tumours grown in Rag1-/-mice treated for 2 weeks with 30 mg/kg RMC-4550 and/or 100 mg/kg RMC-4998. Grey area indicates treatment period. Data are mean tumour volumes ± SEM; n=7-8 mice per group. Analysis was performed using two-way ANOVA. (c) qPCR analysis of immune genes of KPAR^G12C^ orthotopic lung tumours treated for 2 days with 30 mg/kg RMC-4550 and/or 100 mg/kg RMC-4998. Data are mean values ± SD. Each dot represents one tumour, 2 tumours per mouse. Statistics were calculated using one-way ANOVA.

**Supplementary Figure 3.**
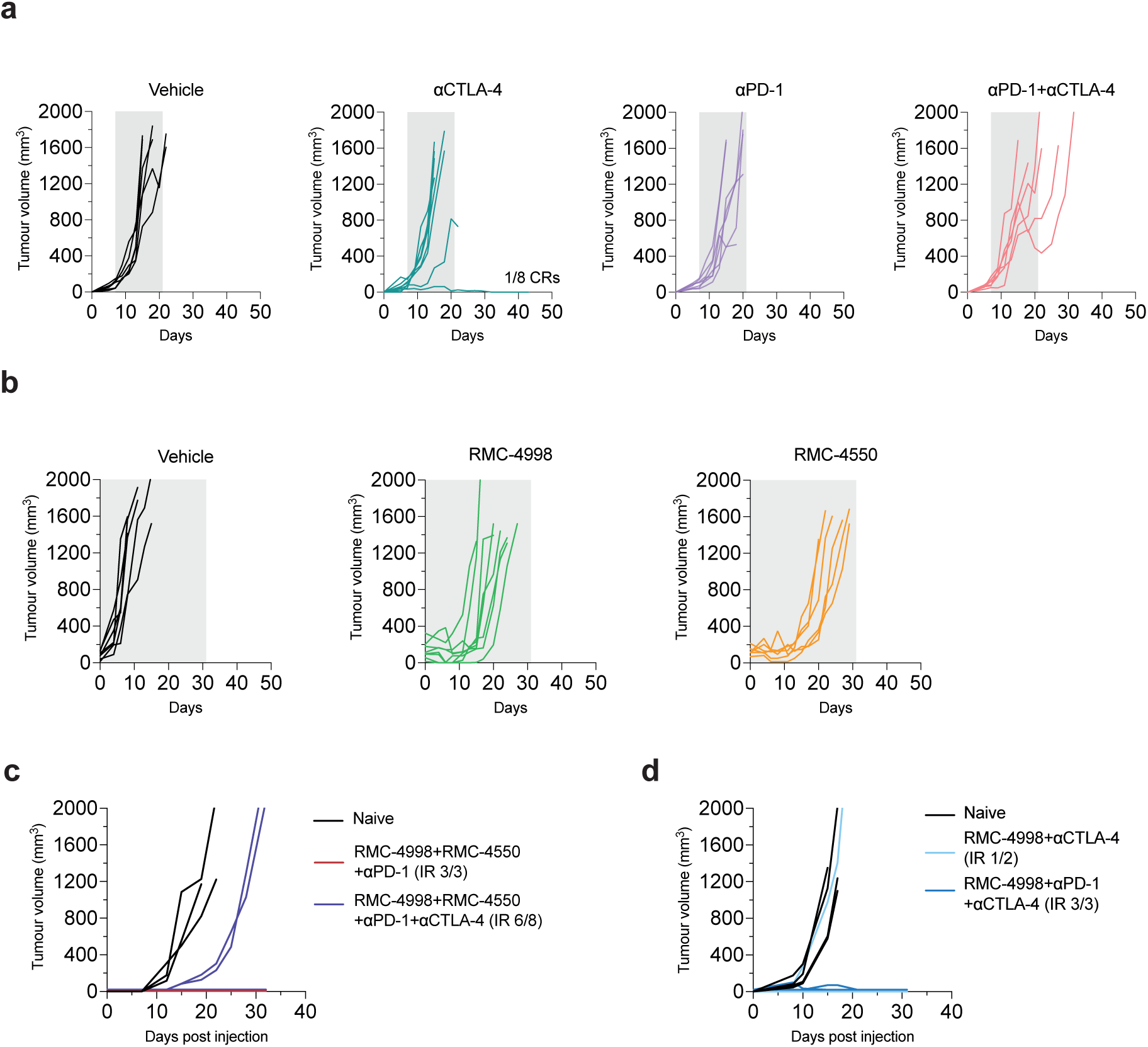
Combination of RMC-4998 and RMC-4550 sensitises 3LL-ΔNRAS subcutaneous tumours to immunotherapies. (a) Individual tumour volumes of mice in Fig. 3a. 3LL-ΔNRAS subcutaneous tumours treated with 10 mg/kg anti-PD-1, 5 mg/kg anti-CTLA-4 or the combination. Grey area indicates treatment period. Number of complete regressions (CR) is indicated. (b) Individual tumour volumes of mice in Fig. 3b. 3LL-ΔNRAS subcutaneous tumours treated with vehicle, 100 mg/kg RMC-4998 or 30 mg/kg RMC-4550. Grey area indicates treatment period. (c-d) Mice in Fig. 3c (c) and Fig. 3d (d) that rejected the primary tumour were rechallenged on the opposite flank and tumour volume was measured. Number of mice that achieved immune rejections (IR) is indicated. Naïve mice of similar age were used as control. Legends indicate the treatment that the primary tumour received.

**Supplementary Figure 4.**
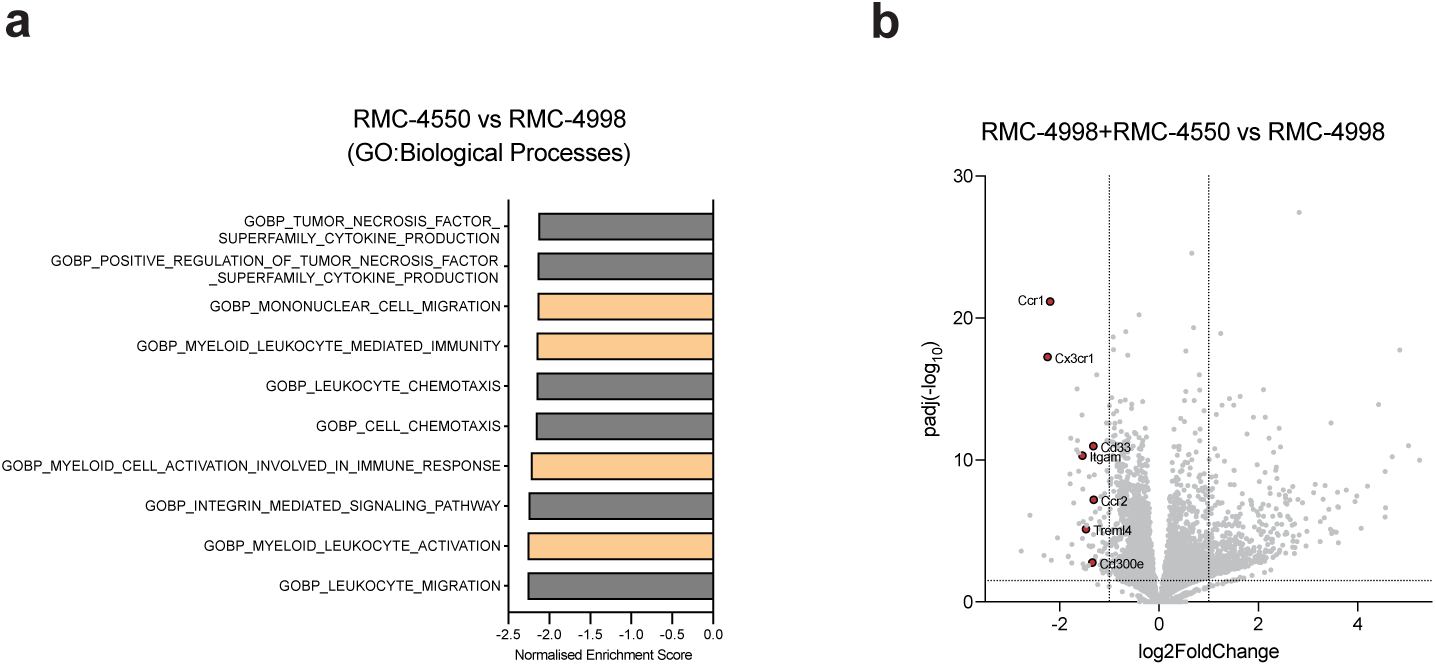
RMC-4988 and RMC-4550 alter the myeloid cell-landscape in the TME of 3LL-ΔNRAS lung tumours. (a) Summary of top 10 significantly (FDR < 0.05) depleted pathways in RMC-4550 treated tumours compared to RMC-4998 treated tumours (MSigDB GO:Biological Processes). (b) Volcano plot highlighting selected significant (p < 0.05) genes from 3LL-ΔNRAS lung tumours treated for 7 days with combination of 100 mg/kg RMC-4998 and 30 mg/kg RMC-4550 vs 100 mg/kg RMC-4998.

**Supplementary Figure 5.**
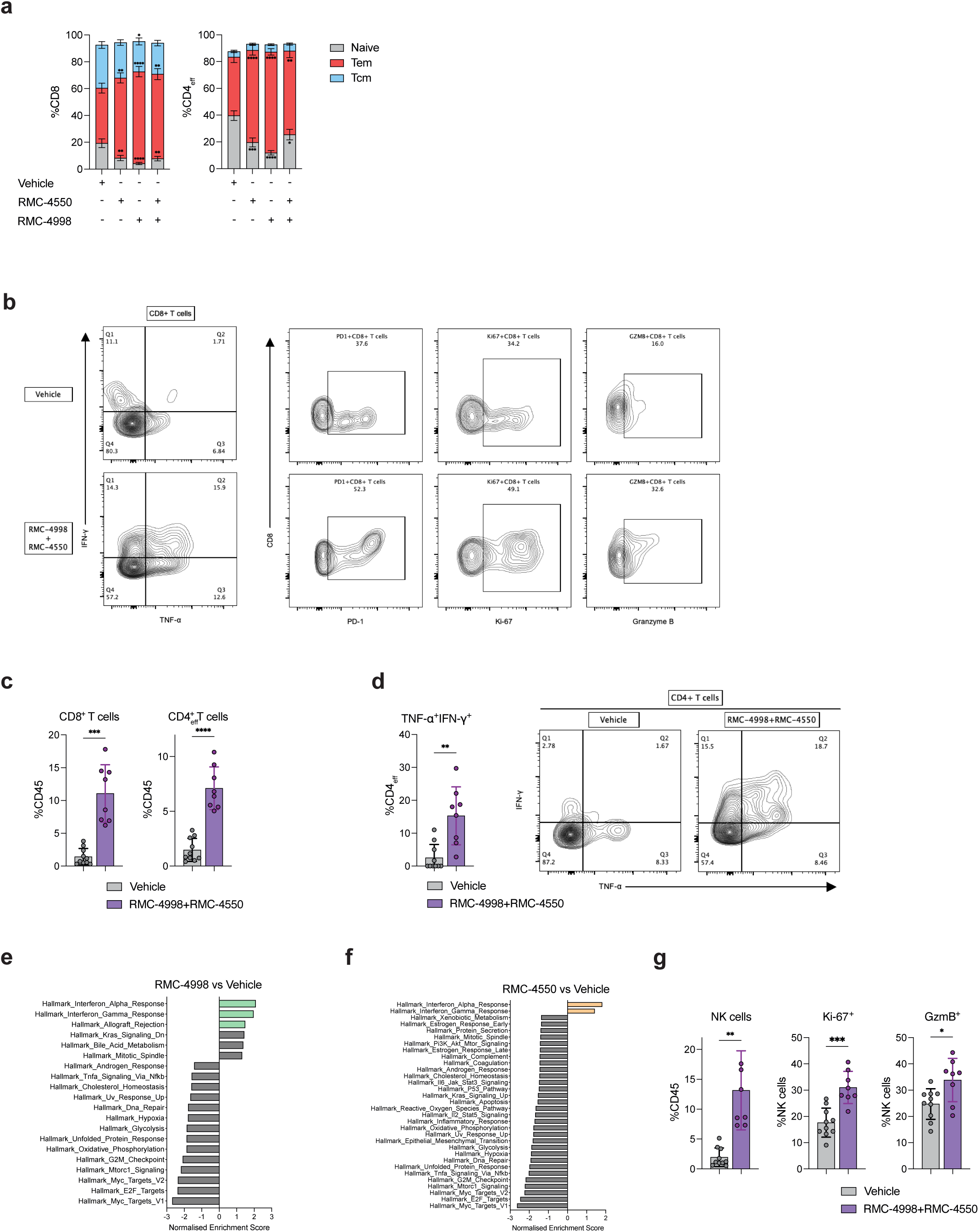
RMC-4988 and RMC-4550 induce pan lymphocytic antitumour response in the TME of 3LL-ΔNRAS lung tumours. (a) Frequency of Naïve (CD44^-^CD62L^+^), Effector memory (CD44^+^CD62L^-^) and Central memory (CD44^+^CD62L^+^) CD8^+^ and CD4^+^ T cells from Fig. 4g. Data ± SEM of two independent experiments. (b) Representative flow plots of PD-1^+^, Ki-67^+^, TNF-α^+^IFN-γ^+^, GzmB^+^ CD8^+^ T cells from Fig. 5e. (c, d) Frequency of CD8^+^ and CD4^+^ T cells from Fig. 5e (c) and frequency of TNF-α^+^IFN-γ^+^ CD4^+^ T cells (d) along with representative flow plots. Data are mean values ± SD. Each dot represents a mouse. Analysis was performed using two-tailed Student’s t-test. (d, e) Summary of significantly (FDR < 0.05) down- or upregulated pathways in tumours treated with 100 mg/kg RMC-4998 (d) or 30 mg/kg RMC-4550 (e) compared vehicle treated tumours (MSigDB Hallmarks). (g) Frequency of NK cells, Ki-67^+^ and GzmB^+^ NK cells in 3LL-ΔNRAS lung tumours treated for 7 days with combination of 100 mg/kg RMC-4998 and 30 mg/kg RMC-4550. Data are mean values ± SD. Each dot represents one mouse. Analysis was performed using two-tailed Student’s t-test.

**Supplementary Figure 6.**
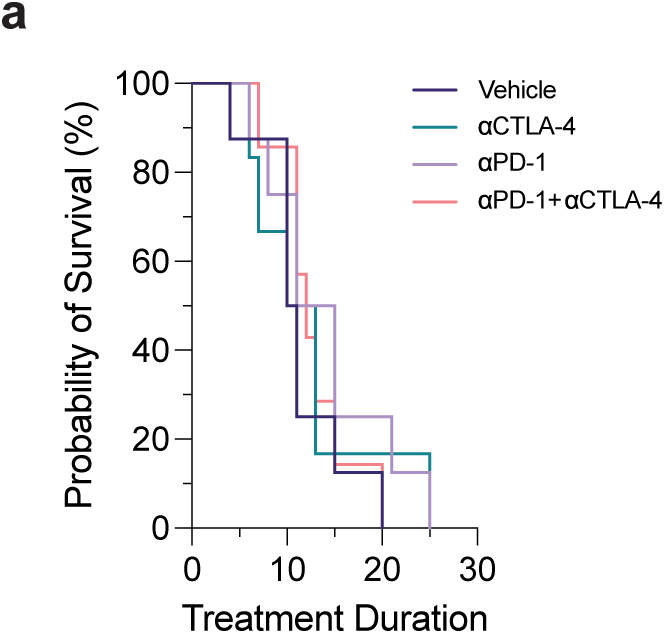
RMC-4998 and RMC-4550 synergise with ICB an orthotopic immune-excluded anti-PD-1 resistant model of NSCLC. (a) Survival of mice bearing 3LL-ΔNRAS orthotopic lung tumours treated with 4 doses of 10 mg/kg anti-PD-1 or 5 mg/kg anti-CTLA-4 or their combination within 2 weeks (n=6-8).

**Supplementary Figure 7.**
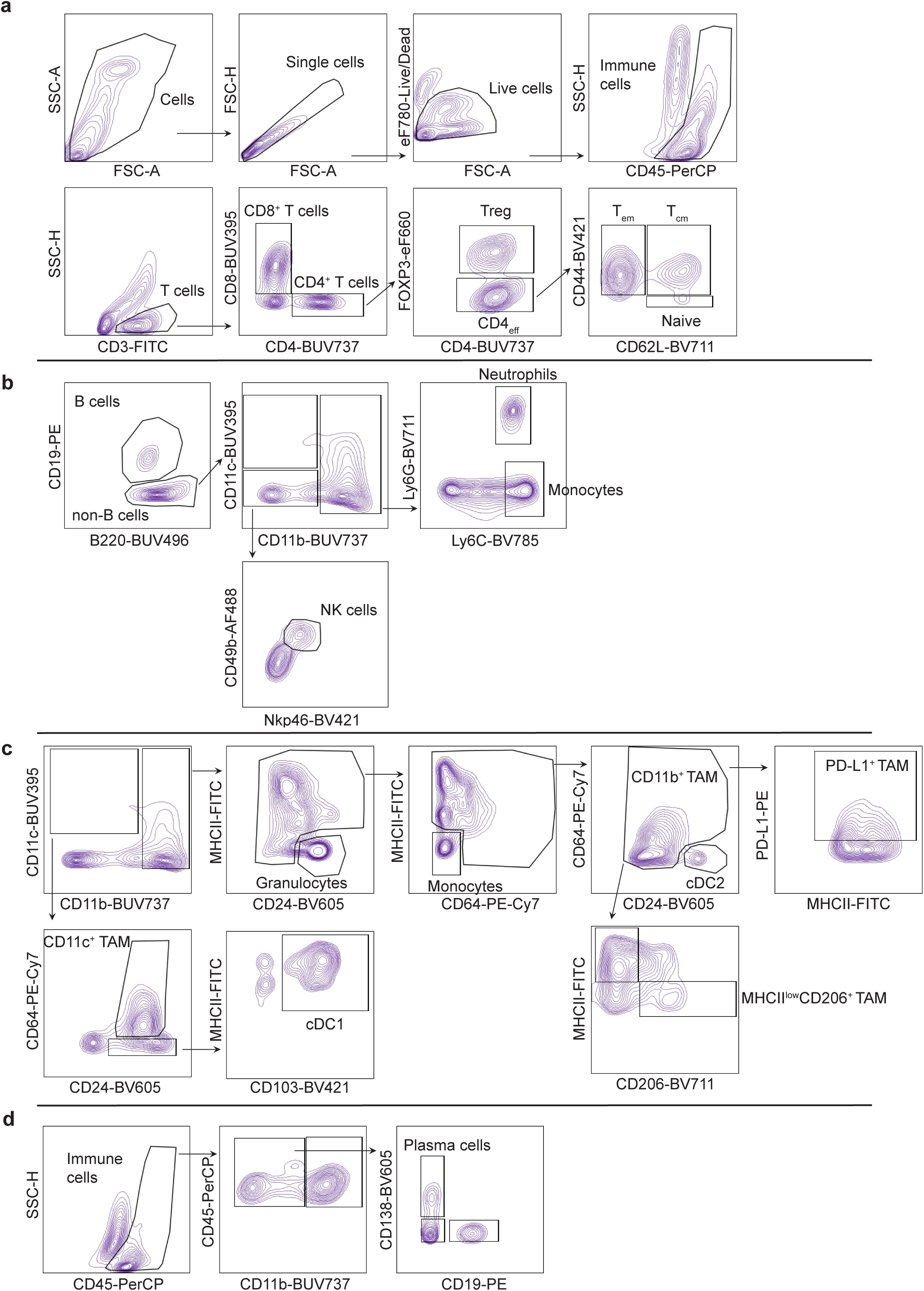
Flow cytometry gating strategies. (a) Gating flow strategy for acquiring single, live, CD45^+^ cells and further gating of T cells populations. (b-d) Gating flow strategy of CD45^+^ cells to acquire B cells, monocytes, neutrophils and NK cells (b), CD11b^+^ TAMs, CD11c^+^ TAMs, cDC1s and cDC2s (c) and plasma cells (d).

**Supplementary Information Table 1.**
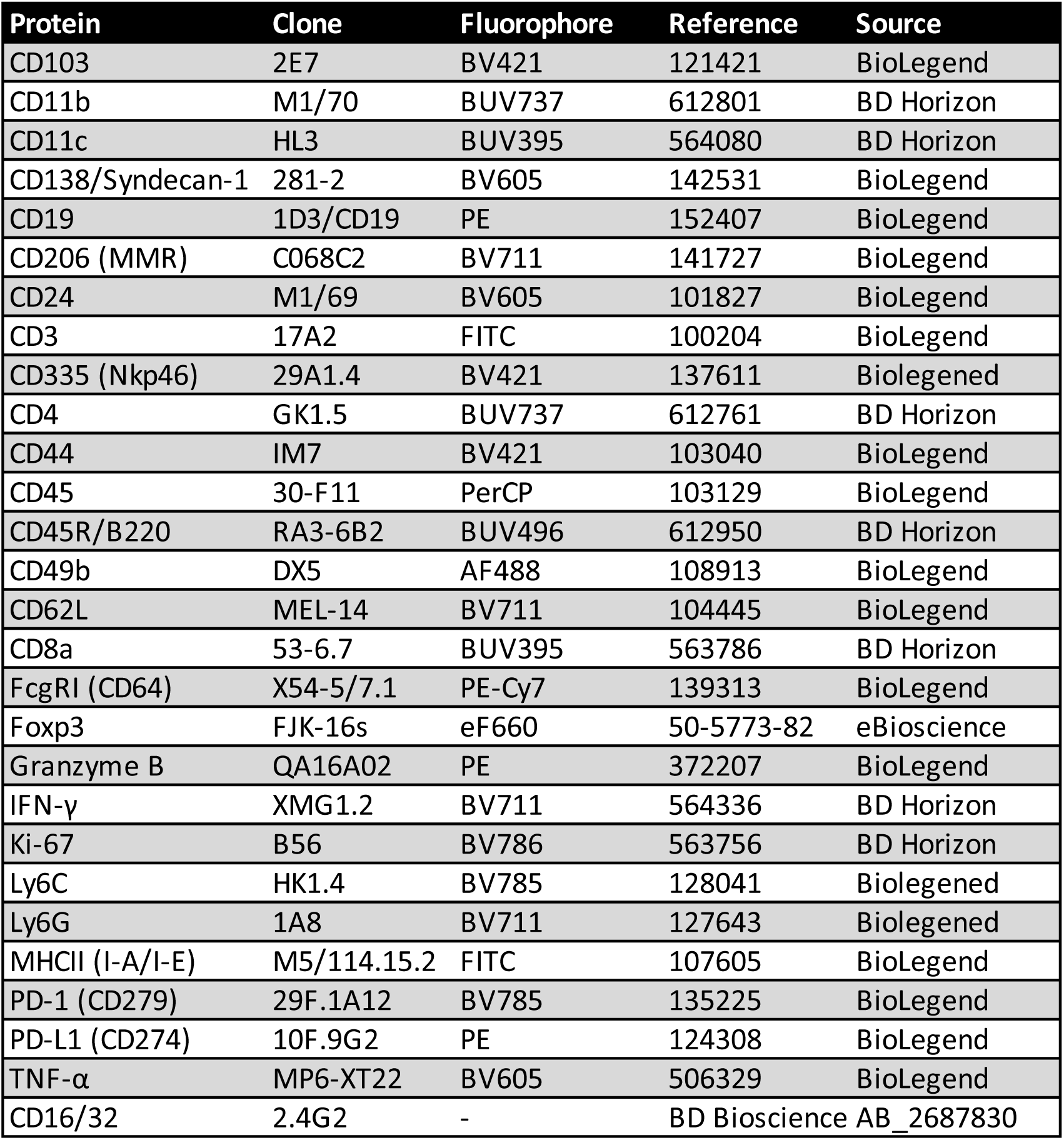
Flow cytometry antibodies.

**Supplementary Information Table 2.** Western blot antibodies.

**Supplementary Information Table 1. Flow cytometry antibodies**
| Protein | Clone | Fluorophore | Reference | Source |
| --- | --- | --- | --- | --- |
| Anti-S6 | 54D2 | - | 2238583 | Cell Signalling |
| Anti-p-S6 (Ser235/236) | Polyclonal | - | 331679 | Cell Signalling |
| Anti-Erk1/2 | 3A7 | - | 10695739 | Cell signalling |
| Anti-p-Erk1/2 (Thr202/Tyr204) | 9101 | - | 331646 | Cell Signalling |
| Anti-Akt | 40D4 | - | 1147620 | Cell Signalling |
| Anti-p-Akt (Ser473) | D9E | - | 2315049 | Cell Signalling |
| Anti-Vinculin | VIN-11-5 | - | 2877646 | Sigma |

## References

1. Prior, I. A., Hood, F. E. & Hartley, J. L. The frequency of ras mutations in cancer. Cancer Res 80, 2969–2974, (2020).

2. Bray, F. et al. Global cancer statistics 2018: GLOBOCAN estimates of incidence and mortality worldwide for 36 cancers in 185 countries. CA Cancer J Clin 68, 394–424 (2018).

3. Simanshu, D. K., Nissley, D. V. & McCormick, F. RAS Proteins and Their Regulators in Human Disease. Cell vol. 170, 17–33, (2017).

4. Yang, H., Liang, S. Q., Schmid, R. A. & Peng, R. W. New horizons in KRAS-mutant lung cancer: Dawn ager darkness. Fron2ers in Oncology vol. 9:953, (2019).

5. Hunter, J. C. et al. Biochemical and structural analysis of common cancer-associated KRAS mutations. Molecular Cancer Research 13, 1325–1335 (2015).

6. Hallin, J. et al. The KRASG12C inhibitor MRTX849 provides insight toward therapeutic susceptibility of KRAS-mutant cancers in mouse models and patients. Cancer Discov 10, 54–71, (2020).

7. Lanman, B. A. et al. Discovery of a Covalent Inhibitor of KRASG12C (AMG 510) for the Treatment of Solid Tumors. J Med Chem 63, 52–65, (2020).

8. Jänne, P. A. et al. Adagrasib in Non–Small-Cell Lung Cancer Harboring a KRAS G12C Mutation . New England Journal of Medicine 387, 120–131, (2022).

9. Skoulidis, F. et al. Sotorasib for Lung Cancers with KRAS p.G12C Mutation. New England Journal of Medicine 384, 2371–2381, (2021).

10. de Langen, A. J. et al. Sotorasib versus docetaxel for previously treated non-small-cell lung cancer with KRASG12C mutation: a randomised, open-label, phase 3 trial. The Lancet 401, 733–746, (2023).

11. Awad, M. M. et al. Acquired Resistance to KRAS G12C Inhibition in Cancer . New England Journal of Medicine 384, 2381–2393, (2021).

12. Zhao, Y. et al. Diverse alterations associated with resistance to KRAS(G12C) inhibition. Nature 599, 679–683, (2021).

13. Ryan, M. B. et al. Vertical pathway inhibition overcomes adaptive feedback resistance to KrasG12C inhibition. Clinical Cancer Research 26, 1633–1643, (2020).

14. Ryan, M. B. et al. KRASG12C-independent feedback activation of wild-type RAS constrains KRASG12C inhibitor efficacy. Cell Rep 39(12):110993, (2022).

15. Fedele, C. et al. SHP2 inhibition diminishes KRASG12C cycling and promotes tumor microenvironment remodeling. Journal of Experimental Medicine 218(1):e20201414, (2021).

16. Ho, C. S. L. et al. HER2 mediates clinical resistance to the KRASG12C inhibitor sotorasib, which is overcome by co-targeting SHP2. Eur J Cancer 159, 16–23, (2021).

17. Liu, C. et al. Combinations with allosteric SHP2 inhibitor TNO155 to block receptor tyrosine kinase signaling. Clinical Cancer Research 27, 342–354, (2021).

18. Adachi, Y. et al. Epithelial-to-mesenchymal transition is a cause of both intrinsic and acquired resistance to KRAS G12C inhibitor in KRAS G12C-mutant non-small cell lung cancer. Clinical Cancer Research 26, 5962–5973, (2020).

19. Santana-Codina, N. et al. Defining and Targeting Adaptations to Oncogenic KRASG12C Inhibition Using Quantitative Temporal Proteomics. Cell Rep 30, 4584–4599, (2020).

20. Xue, J. Y. et al. Rapid non-uniform adaptation to conformation-specific KRAS(G12C) inhibition. Nature 577, 421–425, (2020).

21. Schulze, C. J. et al. Chemical remodeling of a cellular chaperone to target the active state of mutant KRAS. Science 381, 794–799, (2023).

22. Jänne, P. A. et al. Abstract PR014: Preliminary safety and anK-tumor activity of RMC-6291, a first-in-class, tri-complex KRASG12C(ON) inhibitor, in patients with or without prior KRASG12C(OFF) inhibitor treatment. Mol Cancer Ther 22, PR014–PR014 (2023).

23. Liao, W. et al. KRAS-IRF2 Axis Drives Immune Suppression and Immune Therapy Resistance in Colorectal Cancer. Cancer Cell 35, 559–572, (2019).

24. Mugarza, E. et al. Therapeutic KRASG12C inhibition drives effective interferon-mediated antitumor immunity in immunogenic lung cancers. Sci Adv 8(29):eabm8780, (2022).

25. Canon, J. et al. The clinical KRAS(G12C) inhibitor AMG 510 drives anti-tumour immunity. Nature 575, 217–223, (2019).

26. Boumelha, J. et al. An Immunogenic Model of KRAS-Mutant Lung Cancer Enables Evaluation of Targeted Therapy and Immunotherapy Combinations. Cancer Res 82, 3435–3448, (2022).

27. Gandhi, L. et al. Pembrolizumab plus Chemotherapy in Metastatic Non–Small-Cell Lung Cancer. New England Journal of Medicine 378, 2078–2092, (2018).

28. Reck, M. et al. Pembrolizumab versus Chemotherapy for PD-L1–Positive Non–Small-Cell Lung Cancer. New England Journal of Medicine 375, 1823–1833, (2016).

29. Reck, M., Remon, J. & Hellmann, M. D. First-Line Immunotherapy for Non–Small-Cell Lung Cancer. Journal of Clinical Oncology vol. 40, 586–598, (2022).

30. Quintana, E. et al. Allosteric inhibition of SHP2 stimulates antitumor immunity by transforming the immunosuppressive environment. Cancer Res 80, 2889–2902, (2020).

31. Xu, Z. et al. Endothelial deletion of SHP2 suppresses tumor angiogenesis and promotes vascular normalization. Nat Commun 12(1):6310, (2021).

32. Nichols, R. J. et al. RAS nucleotide cycling underlies the SHP2 phosphatase dependence of mutant BRAF-, NF1- and RAS-driven cancers. Nat Cell Biol 20, 1064–1073, (2018).

33. Sisler, D. J. et al. Evaluation of KRASG12C inhibitor responses in novel murine KRASG12C lung cancer cell line models. Front Oncol 13:1094123, (2023).

34. Zagorulya, M. et al. Tissue-specific abundance of interferon-gamma drives regulatory T cells to restrain DC1-mediated priming of cytotoxic T cells against lung cancer. Immunity 56, 386–405, (2023).

35. Simpson, T. R. et al. Fc-dependent depletion of tumor-infiltrating regulatory t cells co-defines the effcacy of anK-CTLA-4 therapy against melanoma. Journal of Experimental Medicine 210, 1695–1710, (2013).

36. Christofides, A., et al. SHP-2 and PD-1-SHP-2 signaling regulate myeloid cell differentiation and antitumor responses. Nat Immunol 24, 55–68, (2023).

37. Niogret, C. et al. Shp-2 is critical for ERK and metabolic engagement downstream of IL-15 receptor in NK cells. Nat Commun 10:1444, (2019).

38. Marasco, M. et al. Molecular mechanism of SHP2 activation by PD-1 stimulation. Sci Adv 6(5):eaay4458, (2020).

39. Haas, L. et al. Acquired resistance to anti-MAPK targeted therapy confers an immune-evasive tumor microenvironment and cross-resistance to immunotherapy in melanoma. Nat Cancer 2, 693–708, (2021).

40. Ghiringhelli, F. et al. Activation of the NLRP3 inflammasome in dendritic cells induces IL-1Β-dependent adaptive immunity against tumors. Nat Med 15, 1170–1179, (2009).

41. Canton, J. et al. The receptor DNGR-1 signals for phagosomal rupture to promote cross-presentation of dead-cell-associated antigens. Nat Immunol 22, 140–153, (2021).

42. Hu, H. et al. Oncogenic KRAS signaling drives evasion of innate immune surveillance in lung adenocarcinoma by activating CD47. Journal of Clinical Inves2ga2on 133(2):e153470, (2023).

43. Huang, A. C. et al. T-cell invigoration to tumour burden ratio associated with anK-PD-1 response. Nature 545, 60–65, (2017).

44. Xiao, P. et al. Myeloid-restricted ablation of Shp2 restrains melanoma growth by amplifying the reciprocal promotion of CXCL9 and IFN-γ production in tumor microenvironment. Oncogene 37, 5088–5100, (2018).

45. Bill, R. et al. CXCL9:SPP1 macrophage polarity identifies a network of cellular programs that control human cancers. Science 381, 515–524, (2023).

46. Rota, G. et al. Shp-2 Is Dispensable for Establishing T Cell Exhaustion and for PD-1 Signaling In Vivo. Cell Rep 23, 39–49, (2018).

47. Ng, K. W. et al. Antibodies against endogenous retroviruses promote lung cancer immunotherapy. Nature 616, 563–573, (2023).

48. Bod, L. et al. B-cell-specific checkpoint molecules that regulate anti-tumour immunity. Nature 619, 348–356, (2023).

49. Saijo, H., et al. AnK-CTLA-4 Antibody Might Be Effective Against Non-small Cell Lung Cancer With Large Size Tumor. An2cancer Res 43, 4155–4160, (2023).

50. Cascone, T. et al. Neoadjuvant chemotherapy plus nivolumab with or without ipilimumab in operable non-small cell lung cancer: the phase 2 plarorm NEOSTAR trial. Nat Med 29, 593–604, (2023).

51. Yofe, I. et al. Anti-CTLA-4 antibodies drive myeloid activation and reprogram the tumor microenvironment through FcγR engagement and type I interferon signaling. Nat Cancer 3, 1336–1350, (2022).

52. Chour, A., et al. Brief Report: Severe Sotorasib-Related Hepatotoxicity and Non-Liver Adverse Events Associated With Sequential AnK–Programmed Cell Death (Ligand)1 and Sotorasib Therapy in KRASG12C-Mutant Lung Cancer. in Journal of Thoracic Oncology vol. 18, 1408–1415, (2023).

53. Li, B. T. et al. OA03.06 CodeBreaK 100/101: First Report of Safety/Effcacy of Sotorasib in Combination with Pembrolizumab or Atezolizumab in Advanced KRAS p.G12C NSCLC. Journal of Thoracic Oncology 17, (2022).

54. Zaw Thin, M., et al. Micro-CT acquisition and image processing to track and characterize pulmonary nodules in mice. Nat Protoc 18, 990–1015, (2023).

55. Molina-Arcas, M. et al. Development of combination therapies to maximize the impact of KRAS-G12C inhibitors in lung cancer. Sci Transl Med 11(510):eaaw7999, (2019).

